# BRD4 recruitment desilences transcription without erasure or depletion of repressive chromatin

**DOI:** 10.64898/2026.04.10.717856

**Authors:** Christopher J. Brandon, Sarah Robinson-Thiewes, Mangesh Kaulage, Wojciech Rosikiewicz, Matthew J. Cuneo, Joseph Brett, Jindpreet Kandola, Walter H. Lang, Jonathan Low, Ashraf Mohammed, Adithi Danda, Sam Rider, Marcus Valentine, Jason Ochoada, Brandon Young, Theresa Nguyen, Sandra J. Kietlinska, Aaron B. Taylor, Burkhard Hoeckendorf, Patrick Rodrigues, Wenwei Lin, Khaled Khairy, Beisi Xu, Anang A. Shelat, Taosheng Chen, Tanja Mittag, Aseem Z. Ansari

**Author notes:** These authors contributed equally to this work (Christopher J. Brandon, Sarah Robinson-Thiewes, Mangesh Kaulage).

## Abstract

How genes are desilenced within mesoscale repressive chromatin is a crucial yet poorly understood phenomenon. Prevailing models posit that methylation of lysine-9 of histone H3 (H3K9me3) engages heterochromatin protein 1 (HP1) to drive chromatin compaction and transcriptional silencing. The erasure of this repressive mark and its replacement with acetyl/acyl groups recruits positive factors such as BRD4/BET to elicit gene transcription. We report that in Friedreich’s ataxia, a synthetic gene regulator (SynGR1) drives transcription across repressive chromatin *without* removal or replacement of H3K9me3 or HP1. By selectively recruiting BRD4/BET into repressive GAA-repeats in frataxin (*FXN*), SynGR1 creates a paradoxical state where gene transcription and repressive chromatin co-exist. Contrary to convention, we find that BRD4 readily partitions into phase separated HP1 condensates in vitro and into HP1 puncta in patient-derived cells, thus presenting a mechanistic explanation for desilencing transcription without the dispersal of mesoscale repressive chromatin. Epigenetic drugs that gate sequential steps in transcription, synergistically stimulate *FXN* expression while concomitantly increasing, rather than eliminating, repressive H3K9me3 and HP1 levels. More broadly, this study highlights the dynamic nature of repressive chromatin and the context-dependence of epigenetic marks in regulating gene expression.

## Main

Over half of the human genome is composed of repetitive sequence elements that are ensconced in repressive heterochromatin^1^. A signature mark of repressive heterochromatin at repeat elements is the trimethylated lysine 9 of histone 3 (H3K9me3)^2,3^. In addition to heterochromatin at pericentromeric repeats, H3K9me3 is also found at genes that are repressed in specific cell lineages, and even in some regions of euchromatin^4–10^. While the role of H3K9me3 in euchromatin remains a mystery, the enrichment of this mark strongly correlates with gene silencing^6–8^. Consistent with key roles in regulating gene expression, aberrant patterning of heterochromatin marks is linked to multiple human diseases^2^.

In Friedreich’s ataxia (FRDA/FA), a life-limiting neurodegenerative disease, expression of frataxin (*FXN*) is downregulated by pathogenic hyper-expansion of GAA trinucleotide repeats within the first intron of the *FXN* gene^11,12^. This repeat expansion is accompanied by increased deposition of H3K9me3 mark^13–18^. The extent of GAA repeat expansions correlates positively with H3K9me3 levels and inversely with gene transcription, demonstrating a functional role for this signature repressive mark in silencing *FXN* expression^17^. Placing expanded GAA repeats at unrelated genomic loci^13^ or downstream of heterologous gene promoters^15^ results in increased H3K9me3 deposition at the new locus. Conversely, augmenting the levels of active acetyl-lysine marks via histone deacetylase (HDAC) inhibitors alleviates the repressive impact of H3K9me3 on *FXN* in patient-derived cell types and in animal models of the disease^14,18–20^.

We previously developed a class of synthetic gene regulators (SynGRs) to restore *FXN* expression in patient-derived cells^21–23^. The prototype SynGR1 (formerly named SynTEF1) is a hetero-bifunctional molecule composed of a GAA-repeat binding polyamide (PA1)^23,24^ tethered to JQ1, a small molecule that binds the BET family of proteins (Fig. S1A). BET proteins, including the well-studied member BRD4, interact with acylated lysine residues on histone tails and recruit the transcriptional machinery^25^. SynGR1 selectively binds the pathogenic GAA repeats, recruits BRD4/BET to the repressive chromatin, and licenses RNA polymerase II (Pol II) transcriptional elongation across the repressive GAA repeats^21^.

Here, we utilized SynGR1 as a precision-tailored mechanistic tool to test conventional models of gene regulation which postulate that desilencing a repressed gene should result in erasure of H3K9me3 marks and removal of HP1 proteins (Fig. 1a). Instead, we found that *FXN* expression not only failed to erode this repressive mark, but it led to an increase in H3K9me3 and HP1 levels. Furthermore, contrary to current models that HP1 and BRD4 form mutually incompatible phase separated condensates, we found that BRD4 partitions into HP1-DNA condensates at physiological concentrations. SynGR1 leverages this naturally occurring cellular phenomenon to enrich BRD4 at repressive GAA-repeats in reconstituted systems in vitro and at *FXN* in FRDA cells. Taken together, these results challenge conventional paradigms and suggest a parsimonious mechanism by which transcription is enabled within repressive chromatin.

**Figure 1.**
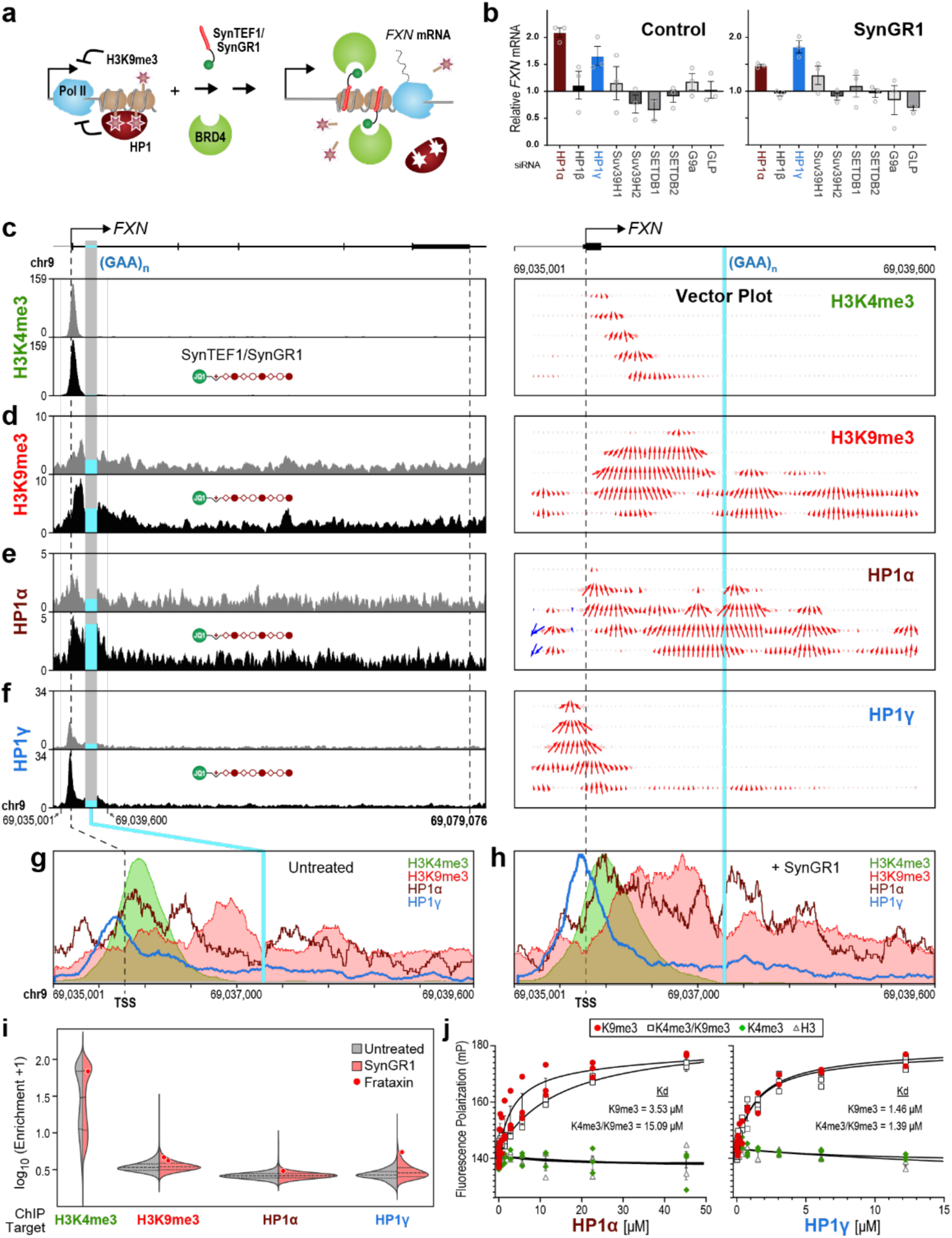
Repressive chromatin marks and HP1 paralogs increase in response to *FXN* gene expression. **(a)** Current models of transcription posit that active transcription leads to removal of repressive histone marks and their readers, such as HP1 proteins. **(b)** Knockdown of specific genes using siRNA on SynGR1-mediated *FXN* expression in FRDA cells. (**c**) Input normalized ChIP-seq and pseudo-velocity vector analysis of H3K4me3 before and after treatment with SynGR1 (n=2; p=n.s.). Psuedo-velocity vector plot stratifies 50bp bins into 5 layers and compares peak changes after treatment to control peaks. The H3K4me3 vector plot visualizes a 3’ movement into the gene body. **(d-f)** ChIP-seq and pseudo-velocity vector plot for H3K9me3 (**d**; n=3; p<1×10^-5^), HP1α (**e**; n=3; p<0.01), and HP1γ (**f**; n=2; p<0.05). **(g-h)** Peak overlay at *FXN* for control (**g**) or SynGR1-treated (**h**) cells which shows changes in peak enrichment after treatment. **(i)** Violin plot of ChIP signals across the genome with or without SynGR1 treatment (red and grey, respectively). FXN enrichment is denoted with red/grey points. **(j)** Fluorescence polarization assays using synthetic 24 amino-acid H3 tail peptides conjugated with fluorescein at the C-terminus. Plots for H3K9me3, H3K4me3, bivalent H3K4me3+K9me3, and unmodified H3 (1-24aa) are displayed. Paired t-tests were performed and showed significance for HP1α (p<0.05) and HP1γ (P<0.001). The dissociation constants (K_d_) were calculated using non-linear least squares curve fit are displayed.

### Persistence of H3K9me3 at actively transcribed *FXN*

While H3K9me3 is predominantly linked to transcriptional repression, this mark infrequently occurs in transcribed regions of the genome^6–9^. To determine if H3K9me3 and HP1 paralogs, modulate *FXN* expression, we knocked down the expression of each of the three HP1 paralogs (α, β, γ) as well as the key enzymes known to methylate H3K9. Consistent with a repressive function, siRNA-mediated knockdown of HP1α and HP1γ paralogs increased *FXN* expression (Fig. 1b, Extended Data Fig. 1h-j). In further support of a repressive role for H3K9me3, the knockdown of Suv39H1, which tri-methylates H3K9, increased expression of *FXN* in FRDA cells (Fig. 1b).

To investigate the rewiring of chromatin landscape in response to *FXN* expression, we monitored ChIP-seq profiles of H3K4me3 and H3K9me3, signature marks of active and repressive chromatin, respectively (Fig. 1c,d). In FRDA lymphocytes (GM15850), the input-normalized H3K4me3 profile shifted subtly downstream, consistent with SynGR1-licensed transcription of *FXN*. A computed pseudo-velocity plot highlights the 3’ vectorial displacement of H3K4me3 enrichment (Fig. 1c, right panel, and Extended Data Figs. 1 and 2). However, to our surprise, the repressive H3K9me3 also increased upon active transcription of *FXN* (Fig. 1d). While uncommon, H3K9me3 marks do occur at some transcribed genes^6–10,26^. Rather than the mark itself, binding and compaction of chromatin by HP1 functions as a barrier to transcribing polymerases^27–31^. Consistent with their repressive effects, HP1 paralogs were enriched at the silenced *FXN* gene in FRDA cells (Fig. 1e, f and Extended Data Fig. 1)^13,32^.

**Figure 2.**
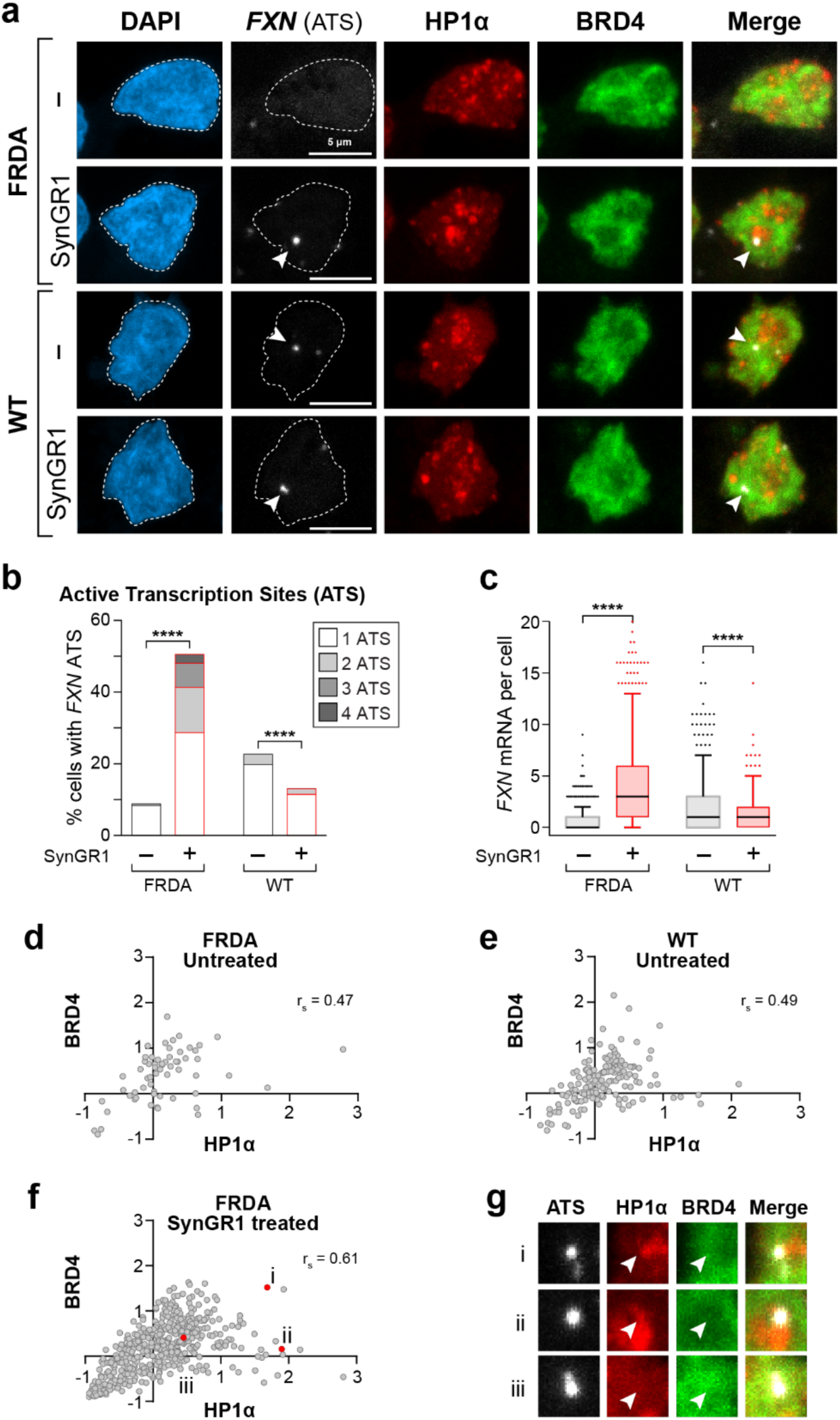
SynGR1 activates *FXN* transcription across the cell population. **(a)** *FXN* RNAscope multiplexed with BRD4 and HP1α immunofluorescence. White arrow, *FXN* active transcription site (ATS); white dashed line, nuclear outline. Scale bar, 5 µm. **(b)** Percentage of cells with a *FXN* ATS in FRDA and WT lymphocytes. ****, p < 0.00001. Number of cells per condition: FRDA SynGR1 = 1147; FRDA untreated = 657; WT SynGR1 = 437; WT untreated = 543. **(c)** Distribution of *FXN* mRNA per cell in FRDA and WT cells. Box plot of 95% confidence interval. ****, p<0.00001. **(d-f)** Correlation between BRD4 and HP1α enrichment at the *FXN* ATS in FRDA untreated (**d**), WT untreated (**e**), and FRDA SynGR1-treated cells (**f**). Spearman’s correlation test used. Number of *FXN* ATS measured: FRDA SynGR1 = 494; FRDA untreated = 61; WT untreated = 166. WT SynGR1 plot (n = 64) is in Extended Data Fig.5. **(g)** HP1α and BRD4 immunofluorescence at the *FXN* ATS (white arrow). Examples i-iii are selected from FRDA SynGR1-treated cells shown in (**f**).

### HP1 enrichment accompanies increased *FXN* expression

Polyamides, the DNA-binding components of SynGRs, are reported to displace HP1 from chromatin^33^. Therefore, we examined if SynGR1 desilences *FXN*, in part, by displacing HP1 proteins from GAA repeats. Unexpectedly, we observed increase in HP1 levels upon *FXN* expression, with HP1α, the most repressive paralog, enriching directly along the path of the elongating Pol II (Fig. 1e). HP1γ, which permits transcription elongation when bound within coding regions^7,34^, displayed increased enrichment at the promoter reflecting its contribution to *FXN* silencing (Fig. 1f,b). To account for the known variability in immunoprecipitation efficiencies of different antibodies^35,36^, we overlaid internally scaled ChIP-seq profiles and observed that an increase in repressive chromatin features accompanied *FXN* desilencing (Fig. 1g, h). Consistent with this observation, input-normalized ChIP signal enrichment of H3K9me3 and HP1α, γ paralogs at desilenced *FXN* were in the top decile of the genome-wide signal distribution for each of three repressive marks (Fig. 1i).

Given the overlap between active (H3K4me3) and repressive (H3K9me3) marks at the *FXN* transcription start site, we measured the ability of the repressive HP1 paralogs to bind histone tails bearing these two opposing marks. Fluorescence polarization was used to quantify binding of histone H3 peptides to purified full length HP1α and HP1γ proteins (Fig. 1j and Extended Data Figs. 3 and 4). Both HP1 paralogs displayed high affinity for di- and tri-methylated lysine-9 (H3K9me2/3) peptides but not for the unmodified (H3), mono/tri methylated lysine-4 (H3K4me1/3), or the phospho-serine-10 (H3S10P) modified peptides that are known to block HP1 binding to nucleosomes (Fig. 1j and Extended Data Fig. 4). Importantly, HP1α and HP1γ bound peptides bearing both opposing K4me3/K9me3 marks, consistent with their ChIP-seq enrichment at regions of *FXN* bearing both histone marks (Fig. 1e-h) and with previous NMR-based structural models of HP1 bound to H3K4/H3K9 bis-methylated histone tails^37^.

### SynGR1 desilences *FXN* expression across the cell population

The overlap in opposing regulatory marks detected by ChIP-seq could well be a consequence of averaging signals across cells in a population wherein a subset of “super-responder” cells have active chromatin marks at the desilenced *FXN* locus whereas the non-responding cells of the population bear repressive chromatin marks at the silent gene. Moreover, between the two alleles within a single cell, the actively transcribed allele may accrue active chromatin marks whereas the silent allele bears repressive marks. We used fluorescence microscopy to investigate whether such conflation of signals explains the superimposed chromatin marks observed in ChIP-seq profiles.

To map actively transcribed sites (ATS), we used the single molecule RNAScope approach which utilizes unique probes that hybridize to *FXN* transcripts and provide means to amplify the fluorescent signal at the transcribed allele. Visual inspection revealed that *FXN* transcripts are rarely observed in untreated FRDA cells (Fig. 2a). SynGR1 treatment elicited a pattern of *FXN* expression across the cell population that was indistinguishable from those observed in *FXN* expressing healthy cells (Fig. 2a, lower panel and Extended Data Fig. 5). To quantitatively analyze our imaging data, we developed a custom ImageJ/FIJI image processing pipeline and used MATLAB tools to segment the images and quantify single mRNA molecules in each treatment condition. In FRDA lymphocytes, SynGR1 treatment increased the fraction of cells with desilenced *FXN* ATSs (Fig. 2b) as well as the polyadenylated mRNA (Fig. 2c), consistent with our previously reported bulk RNA measurements^21,22^. Thus, SynGR1 restores *FXN* expression across the population of diseased cells in a manner that mirrors *FXN* expression patterns in healthy cells.

### HP1α and BRD4 co-localize in cells

To determine if the actively transcribed *FXN* locus co-localizes with BRD4 or HP1α, we combined RNAScope with imaging of these two readers of epigenetic marks via immuno-fluorescence (IF). As expected, these functionally orthogonal proteins primarily enrich in non-overlapping regions of the nucleus, however they do overlap at *FXN* ATS (Fig. 2a). Quantifying BRD4 and HP1α levels at individual *FXN* ATSs and, as a control, at equally sized randomly selected locations within the cell revealed significantly higher overlap between the two opposing effectors at the *FXN* ATS (Fig. 2d-g, Extended Data Fig. 5). In SynGR1-treated FRDA cells, far more *FXN* ATSs were detected, accompanied by a greater correlation, r_s_ = 0.61, of co-localized BRD4 and HP1α (Fig. 2f). Moreover, suggestive of a dynamic process, a broad range in the levels of co-localized HP1α and BRD4 was observed in all conditions (Fig. 2d-g). These results indicate that not only can BRD4 gain access to regions of modest HP1α enrichment, but that it can also be recruited into HP1α puncta, often assumed to reflect phase separated condensates (Fig. 2g).

To determine if the observed co-localization of these opposing factors was a unique feature of the *FXN* locus, we quantified the extent of BRD4 and HP1α co-occurrence across the entire nucleus in WT and FRDA cells. Indicative of a widely occurring cellular phenomenon, low level of co-localization of BRD4 and HP1α was evident under all conditions in WT and FRDA cells (Extended Data Fig. 6). We then quantified BRD4 levels within all HP1α-enriched puncta and normalized the values to the mean signal across the nucleus (i.e., non-punctate nucleoplasm; see Methods). Contrary to conventional models^38,39^, even within dense HP1α puncta, the two functionally orthogonal proteins were not strictly anticorrelated (Extended Data Fig. 6b-d). In other words, our data reveal a naturally occurring cellular phenomenon where the two functionally opposed effector proteins, BRD4 and HP1α, dynamically sample the orthogonal chromatin states. In particular, the data provide single cell and single locus evidence for the co-localization of BRD4 and HP1α at the desilenced *FXN* locus. Our observations are further supported by a recent report of chemically induced dimerization of sfGFP-fused BRD4 and Halo-tagged HP1α in cells^40^.

### BRD4 partitions into reconstituted HP1α condensates

HP1α and BRD4 form functionally and physically distinct condensates and their ability to access orthogonal condensates at physiological concentrations was unexpected^38,39,41^. To determine if this phenomenon can be recapitulated with purified components, we examined if HP1α forms condensates with GAA repeat DNA and whether such condensates permit access to BRD4 without complete or partial dispersion. To enable comparisons with prior studies, we used a 147-base pair (bp) DNA fragment bearing 49 tandem GAA repeats to generate HP1α-DNA condensates. When compared with the well-characterized 601 DNA sequence of identical length^42^, HP1α phase-separated with GAA repeats at comparable saturation concentrations and formed condensates of similar dimensions (Fig. 3a-c). Next, to examine if SynGR1 can access cognate binding sites in HP1α-DNA condensates, we synthesized a (GAA)_3_-selective polyamide (PA1) conjugated to JF646 (Fig. 3d and Extended Data Fig. 7b,c). Consistent with its specificity for GAA repeats^23,24^, PA1-JF646 enriched within HP1α condensates bearing 49 tandem GAA-repeats but not in condensates formed with identically sized 601 DNA, which lacks high affinity PA1/SynGR1 binding sites (Fig. 3e-g). PA1 or PA1-JF646 did not disperse the (GAA)_49_-HP1α condensates even at saturating concentrations (Fig. 3e-h). More importantly, SynGR1 facilitated the partitioning of BRD4 into these condensates at low nanomolar concentrations (Fig. 3i,j) and it did so within seconds (Fig. 3k and Extended Data Fig. 7e). At higher concentrations (>250 nM), BRD4 accessed HP1α condensates without the assistance of SynGR1 (Fig. 3j). This result is consistent with cellular immunofluorescence data above where low levels of BRD4 were observed within HP1α puncta both in healthy and FRDA cells (Extended Data Fig. 6). These results further support the interpretation that BRD4 can access HP1 condensates at some level and that this is a naturally occurring cellular phenomenon which is not unique to pathogenic GAA-repeats and occurs independently of SynGR1.

**Figure 3.**
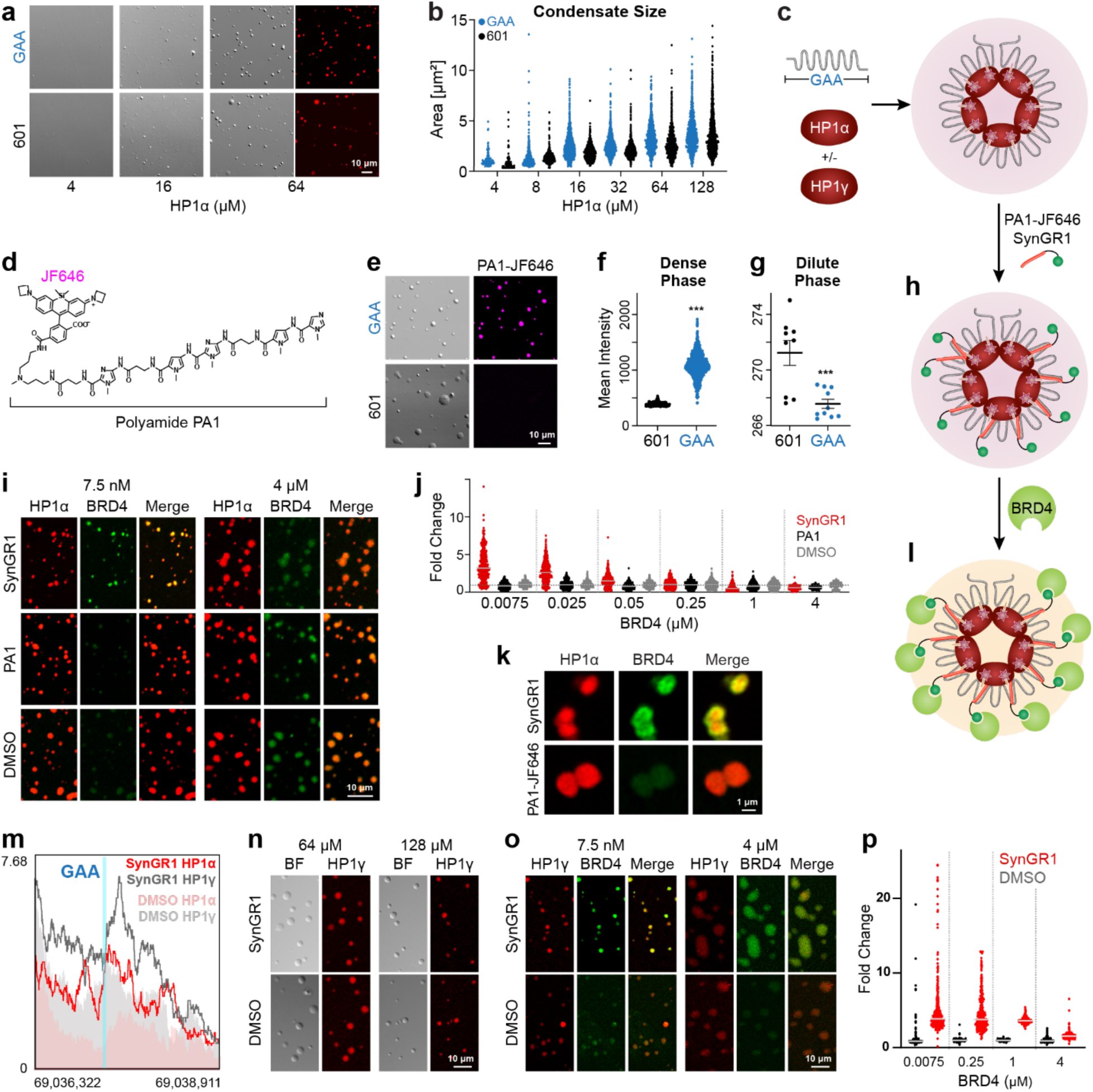
BRD4 can access HP1 condensates. **(a)** Titration of 601 DNA and GAA-repeat DNA with increasing concentrations of HP1α. Samples imaged by DIC microscopy. Fluorescence micrographs of samples at 64 μM HP1α show its enrichment in condensates. **(b)** Condensate area over 28,000 total droplets and 4 replicates. **(c)** Schematic showing HP1α can form condensates with GAA-repeat DNA. **(d)** Chemical structure of PA1-JF646, a polyamide conjugated to fluorophore JF646. **(e)** PA1-JF646 enters HP1α condensates formed with GAA-repeat DNA, but not 601 DNA. **(F)** PA1-JF646 fluorescence intensity in condensates formed with GAA-repeat DNA or 601 DNA. **(g)** Depletion of PA1-JF646 fluorescence intensity from the dilute phase in presence of HP1α-DNA condensates. **(h)** Schematic showing PA1-JF646 and SynGR1 into HP1α-GAA DNA condensates. **(i)** Access to BRD4 into HP1α-GAA DNA condensates in the presence of SynGR1, PA1 or DMSO. **(j)** Quantification of (i) over 15,000 droplets and 2 replicate samples. Fold change is fluorescence intensity normalized to the average intensity in condensates treated with DMSO. **(k)** Super-resolution micrographs showing SynGR1-dependent entry of BRD4 into pre-formed HP1α-DNA condensates. **(l)** Schematic showing SynGR1-mediated BRD4 recruitment to HP1α condensates. **(m)** ChIP-seq profiles of HP1α and HP1γ across GAA repeats with or without SynGR1 treatment. **(n)** Co-condensate formation by 1:1 or 2:1 molar ratio of HP1γ and HP1α with the 49 GAA repeat-bearing DNA. A fraction of HP1γ was fluorescently labeled for imaging. **(o)** SynGR1 helps partition BRD4 into these mixed condensates at 7.5 nM but is not required at higher concentrations of BRD4. **(p)** Quantification of SynGR1-aided partitioning of BRD4 into mixed HP1 condensates.

**Figure 4.**
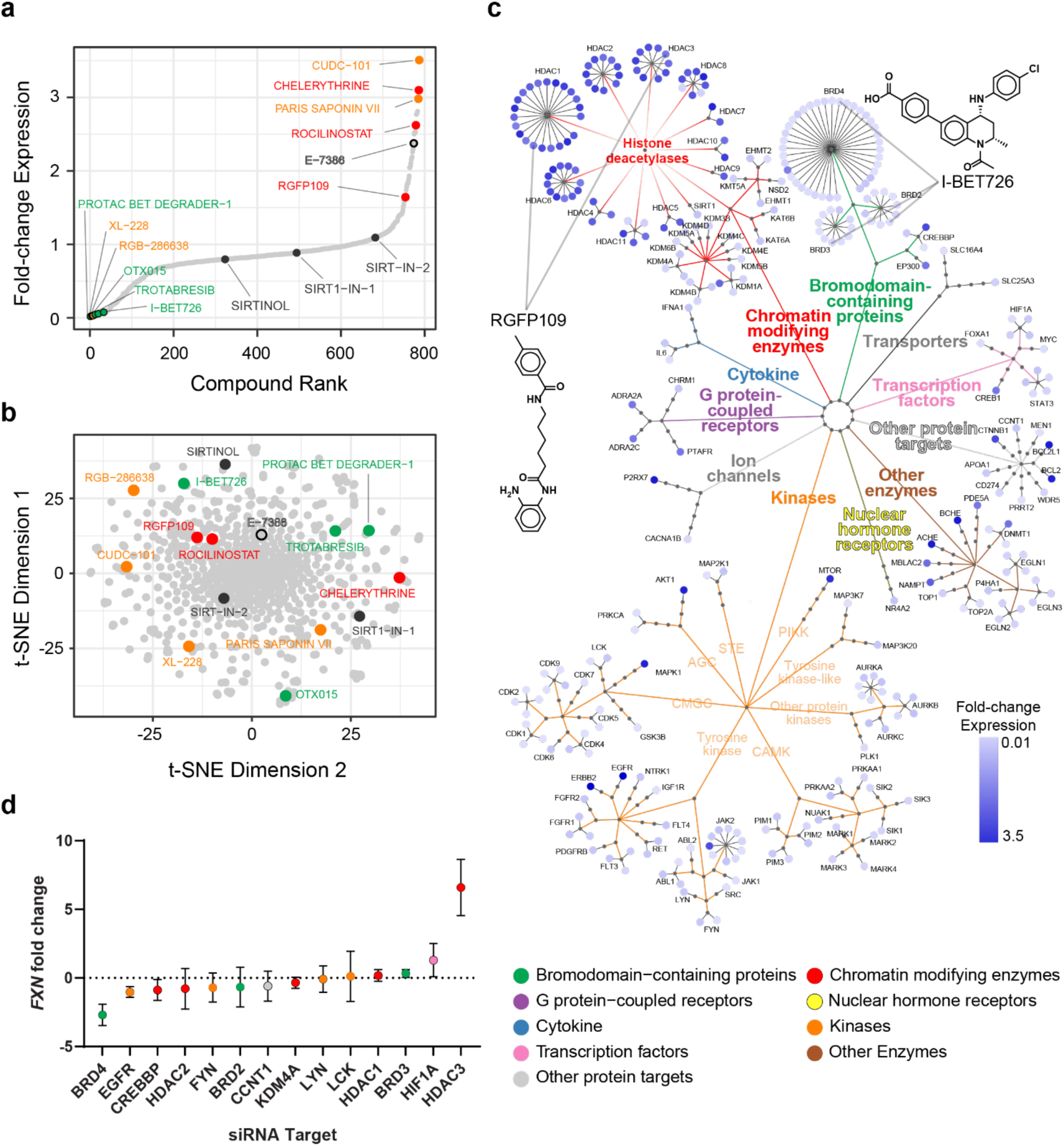
Identification of epigenetic modulators of SynGR1. **(a)** Scatterplot of the fold-change in *FXN* expression induced by combining each member of a library of 791 small molecule epigenetic modulators with SynGR1 compared to SynGR1 alone. Exemplar molecules enhancing, inhibiting, or not changing gene expression relative to SynGR1 alone are colored by compound class. (**b**) Chemical space representation of the molecules from (a) based on projecting the first and second t-SNE dimensions derived from the Morgan molecular fingerprint distance matrix. Exemplar molecules are colored as in (a). **(c)** Activity based hierarchical network graph representing 145 compounds from (a). Molecules are represented as leaf nodes and are color coded by the fold-change in *FXN* expression as reported in (a). **(d)** Plot of the fold-change in *FXN* expression resulting from genetic knockdown of gene targets identified from the epigenetic library screen. Genes are color coded according to the mechanistic classes depicted in (d). Values are plotted as the mean and SD.

In addition to HP1α, HP1γ also enriched across the GAA repeats of the desilenced *FXN* (Fig. 3m). Therefore, we tested whether the two HP1 paralogs, either at equimolar (64 µM each) or at 2:1 molar ratio (128 µM HP1γ: 64 µM HP1α) also form co-condensates with the 147-bp GAA repeat DNA. In both conditions, we observed condensate formation (Fig. 3n). As with HP1α-DNA condensates, SynGR1 readily enriched BRD4 in HP1α/γ co-condensates at nanomolar concentrations. Enrichment of BRD4 within these condensates occurred at higher concentrations in the absence of SynGR1 (Fig. 3o,p). These biophysical studies with reconstituted systems further corroborate ChIP-seq and cell imaging studies which independently demonstrate co-localization of these functionally orthogonal proteins in cells, independent of SynGR1 treatment. While initially identified at the desilenced *FXN*, BRD4 naturally partition into most HP1 puncta in healthy and FRDA cells. To further corroborate this observation, we examined whether BRD4 and HP1 co-localize at non-*FXN* genomic loci. Among a plethora of examples, a class of zinc finger (ZNF) genes were particularly noteworthy because they are enriched in overlapping “contradictory” chromatin marks -repressive H3K9me3 and active H3K36me3 ^43^. A striking overlap in ChIP-seq profiles of BRD4 and HP1 was evident at these ZNF genes and at other transcribed genes that are ensconced in repressive chromatin (Extended Data Fig. 7). Thus, the innate ability of BRD4 to access HP1-rich domains more broadly coincides with transcription within mesoscale repressive chromatin.

### Synergistic activation with epigenetic inhibitors

Given that HP1α and HP1γ repress *FXN* in FRDA cells (Fig. 1b), we hypothesized that pharmacological agents that prevent enrichment of these repressive paralogs would enhance *FXN* expression. Because there are no small molecule inhibitors of HP1, we screened a focused library of 791 epigenetic inhibitors from MedChem Express (see Methods). Members of the library span a diverse chemical space, and a few molecules target other proteins in addition to those with epigenetic roles (Fig. 4a). 39 molecules increased and 106 reduced SynGR1-mediated *FXN* expression (Fig. 4b). Validating our approach, BET inhibitors and degraders that prevent BRD4/BET recruitment diminished *FXN* expression. Amongst the diverse set of molecules that enhance *FXN* expression, the vast majority are histone deacetylase inhibitors (Fig. 4c). We cross-validated these results with siRNA-mediated knockdown of known protein targets of active small molecules. Knockdown of BRD4 attenuated *FXN* expression whereas knockdown of Class 1 HDACs enhanced *FXN* expression (Fig. 4d). Many of the HDAC inhibitors were optimized for oncological applications and have unfavorable toxicity profiles, therefore we focused on the brain-penetrant histone deacetylase inhibitor RGFP109/RG2833 which was developed by Repligen to increase

*FXN* expression in FRDA animal models of the disease^20^. RGFP109 selectively inhibits members of class I histone deacetylases and inhibits the erasure of transcriptionally conducive acetyl-lysine-9 (H3K9ac) chromatin marks (Fig. 5a, b). The retention of H3K9ac marks is also expected to block the placement of the repressive H3K9me3 marks.

**Figure 5.**
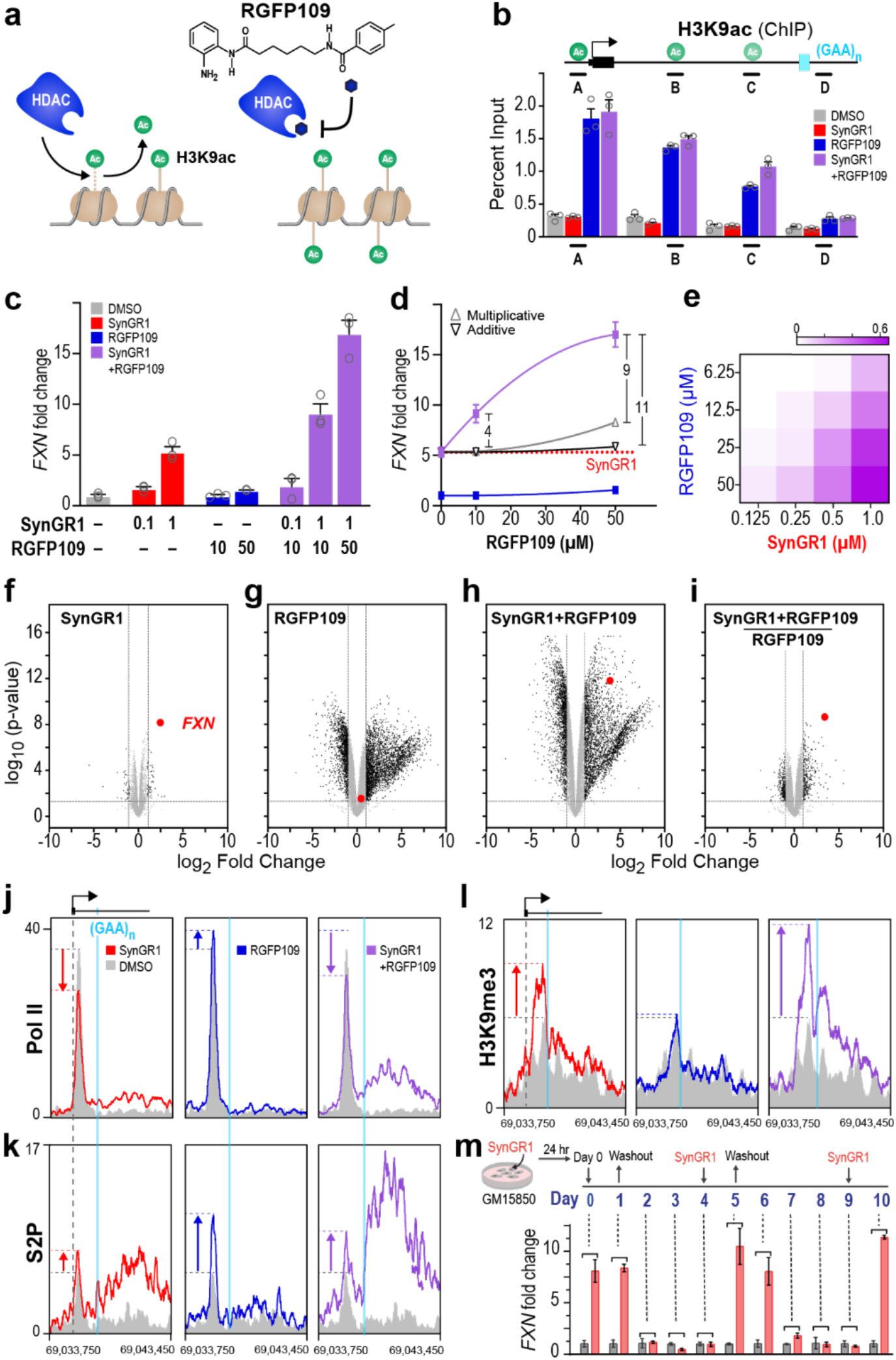
Mechanistic basis for synergistic *FXN* expression. **(a)** Schematic of Histone deacetylase inhibition by RGFP109. **(b)** ChIP-qPCR for H3K9ac normalized to percent input at specified amplicons across the *FXN* gene. **(c)** qRT-PCR analysis of *FXN* expression normalized to DMSO control. Error bars are SEM of 3 replicates. **(d)** Comparative plots of observed *FXN* levels upon single or double agent treatment versus expected additive or multiplicate effects. **(e)** BLISS independence model evaluation of synergy with pairwise combinations of RGFP109 (DMSO, 6.25, 12.5, 25, 50) and SynGR1 (DMSO, 0.125, 0.25, 0.5, and 1 uM) (n=3). **f-i,** Volcano plots comparing RNAseq expression changes in response to SynGR1 **(f)**, RGFP109 **(g)**, SynGR1 + RGFP109 treatment **(h)**, or corrected for RGFP109 in co-treated samples **(i)**. **(j-l),** ChIP-seq profile for Pol II (**j**), phospho-Ser2 (S2P); **(k)**, and H3K9me3 **(l)**, all samples were normalized to input. **(m)**, FRDA patient-derived cells (GM15850) were treated with 1 μM SynGR1 for 24 hrs, washout of SynGR1 was followed by reversion of *FXN* mRNA to repressed levels. Successive retreatment of these *FXN* silenced cells with SynGR1 for 24 hours was followed by washout was use to observe *FXN* mRNA by qPCR. Data represents three independent experiments.

Co-treatment of RGFP109 with SynGR1 showed synergistic *FXN* expression that greatly exceeded the maximal activation achieved by either small molecule on its own (Fig. 5c, d). The BLISS independence model for drug-drug interactions was used to map the synergy landscape of the two effectors of *FXN* expression (Fig. 5e), and the resulting quantitative measure of synergy was consistent with the role of repressive chromatin in attenuating *FXN* expression. Using RNA-seq, we next examined whether the synergistic stimulation of *FXN* by RGFP109 also triggered SynGR1-responsive transcription initiation at other GAA-rich genomic loci. The comprehensive transcriptome-wide profiles identified *FXN* as the primary target of SynGR1 (Fig. 5f), whereas RGFP109 treatment dramatically remodeled the transcriptome, with 2854 transcripts up-regulated and 1366 transcripts down-regulated (Fig. 5g and Extended Data Table 1). The transcriptome-wide perturbation by a freely diffusing HDAC inhibitor is not unexpected, as blocking HDAC activity across the genome is expected to elicit wide-ranging disruptive consequences. In agreement with RT-qPCR results (Fig. 5c, d), the unbiased transcriptomic data showed that co-treatment with SynGR1 and RGFP109 non-linearly raised *FXN* mRNA levels (Fig. 5h). Importantly, when normalized against RGFP109-treated cells, the residual transcriptome profile closely matched that of cells treated with SynGR1 alone (Fig. 5i). These results demonstrate that RGFP109 does not adventitiously enable SynGR1 to stimulate transcription at other genomic loci.

To investigate the basis of synergy between SynGR1 and RGFP109, we measured the levels of transcriptionally conducive H3K9ac marks across *FXN* by quantitative ChIP-qPCR (Fig. 5b). In agreement with previous reports^14^, RGFP109 treatment led to an increase in H3K9ac marks at the promoter (Fig. 5b, amplicon A). However, levels of H3K9ac marks steadily declined upon approach to the GAA repeats. This pattern of H3K9Ac enrichment presents a mechanistic explanation for the observed synergy in the presence of RGFP109 and SynGR1. In untreated FRDA cells, modest levels of H3K9ac marks at the *FXN* promoter would moderate the levels of transcription initiation (Fig. 5b, j). This early regulatory step is not altered by SynGR1, which functions instead by enabling transcriptional elongation by Pol II across the roadblock presented by the repressive GAA repeats downstream. The functional role of SynGR1 in licensing this post-initiation step is evident from the increase in elongating Pol II (S2P/pSer2), downstream of the GAA repeats (Fig. 5k). In contrast, RGFP109 treatment increases H3K9ac levels at the promoter, disrupting promoter-gating and permitting higher levels of transcription initiation^44^ (Fig. 5j, k). Despite higher levels of initiation, only a small fraction of the promoter-loaded Pol II is able to surmount the repressive GAA repeats, resulting in modest gains in *FXN* mRNA levels (Fig. 5k). In the absence of RGFP109, innate promoter gating coupled with SynGR1/BRD4-assisted transcription elongation work together to restore *FXN* expression in FRDA cells to the levels observed in healthy cells. However, in the presence of both small molecules, a larger fraction of the RGFP109-induced Pol II flux through the promoter is further licensed by SynGR1 to transcribe past the repressive repeats, yielding dramatically increased levels of *FXN* transcripts (Fig. 5j, k).

### Over-riding dynamically reinforced H3K9me3 and HP1

Given the high levels of *FXN* expression triggered by the combined action of SynGR1 and RGFP109, we examined if H3K9me3 repressive marks were altered under these extreme conditions. Instead of being eroded or erased as predicted by prevailing paradigms, the levels of H3K9me3 rose substantively over the region spanning the repeats (Fig. 5l). Rather than a dormant epigenetic state, the data revealed that fresh repressive marks are placed in response to active transcription of disease-causing GAA repeats. The hyper-methylation of H3K9 upon co-treatment with RGFP109 and SynGR1 emerges as a consequence of perturbing the balance between dynamic erasure and re-writing of repressive marks.

Reinforcement of repressive marks at transcribed regions is linked to transcript-assisted recruitment of silencing proteins such as Polycomb Repressive Complex (PRC2) or the Human Silencing Hub (HUSH) complex^26,45,46^. To examine if these complexes play a role in repressing *FXN* in FRDA cells, we knocked down multiple members of both complexes and observed modest impact on *FXN* expression (Extended Data Fig. 9). Next, we examined if the increase in repressive chromatin marks led to irreversible silencing of transcription. To test this possibility, we treated FRDA cells with SynGR1 for 24 hours, i.e., conditions under which *FXN* expression triggers an increase in H3K9me3 and HP1 enrichment. After 24 hours, we washed out SynGR1 and over 72 hours monitored the decline in *FXN* mRNA levels to basal levels (Fig. 5m). Once basal levels were reached, we re-treated cells with the same dose and duration of SynGR1 and observed that *FXN* expression was fully restored, indicating that *FXN* transcription had not been irreversibly silenced by the placement of repressive marks. Identical induction and decay profiles were obtained when the cells were treated for the third time. Taken together, the data demonstrate that repressive marks in response to *FXN* expression are dynamic and permit transcription within the context of repressive chromatin.

### Mechanistic and therapeutic implications

HP1α is enriched at repressive GAA-repeats at *FXN* in patient-derived cells. Nonetheless, SynGR1 can access these nuclease inaccessible GAA repeats and partition BRD4 into these repressive HP1α-rich regions. The ability of BRD4 to access HP1-rich puncta in cells and in HP1-DNA condensates in vitro, extends well beyond *FXN* and occurs independently of SynGR1 treatment. This observation necessitates a reconsideration of the widely held assumption that functionally orthogonal BRD4 and HP1α protein do not coacervate. Furthermore, consistent with a functional role, the ability of BRD4/BET inhibitors and degraders, to attenuate *FXN* expression indicates that recruitment of BRD4 within HP1-rich domains suffices to desilence transcription. Extending beyond *FXN*, the co-localization of BRD4 and HP1α/y at numerous other genes that bear contradictory/orthogonal chromatin marks correlates with their finely tuned expression in the context of repressive chromatin.

We note that transcription within H3K9me3-marked regions and recruitment of HP1γ during transcription has been previously reported^7,8^. However, the mechanism by which transcription is enabled without removal of repressive heterochromatin remains poorly understood. Using SynGR1 as a tool to monitor chromatin rewiring as *FXN* is desilenced, provides compelling evidence that recruiting BRD4 suffices to license processive transcription while retaining repressive chromatin state at the transcribed locus. Armed with this mechanistic insight, we evaluated our imaging data and found that, albeit infrequent, BRD4 did co-localize with non-*FXN* HP1α puncta at physiological concentrations. This occurred in a SynGR1-independent manner in both healthy and diseased cells. Moreover, BRD4 co-localizes with HP1 at a wide-array of genes, including a class of zinc finger encoding genes where transcription elongation is greatly attenuated by H3K9me3-enriched repressive chromatin. Collectively, our data reveal a new mechanism by which a gene resident within heterochromatin can be transcribed in response to cellular cues, and instantly re-silenced once the cellular need is met. The ability of SynGR1 to achieve this by selectively recruiting BRD4 presents a parsimonious mechanism that may well be used by transcription factors that enable transcription at silenced genomic loci^47^.

How might BRD4 recruitment achieve a paradoxical state wherein transcription does not cause the erasure of repressive marks or depletion of the effectors of repression? We propose that the rapid association and dissociation kinetics of HP1α for DNA, and the reported viscoelastic properties of the HP1α-DNA condensates^42^, provide a solution to this conundrum. In essence, HP1α-DNA condensates would rebuff high-flux transcription bursts but would permit passage to a slowly elongating Pol II that is assisted by synthetically positioned elongation machinery^42,48^ (Fig. 6). Furthermore, in a negative-feedback loop, the resulting transcripts often recruit repressive epigenetic machinery, such as the HUSH and PRC complex, to silence their own expression^3,49,50^. Similarly, GAA-rich transcripts could help retain HP1 at the transcribed locus and function as a rheostat to moderate the levels of *FXN* expression. By desilencing *FXN* without eroding or erasing the repressive environment imposed at the repeat expansions, we provide more broadly relevant insights into the fluidity and context-dependence of the histone code. While the signature epigenetic marks function as expected by convention, the ease with which they can be overridden to permit expression of critical genes was entirely unexpected.

**Figure 6.**
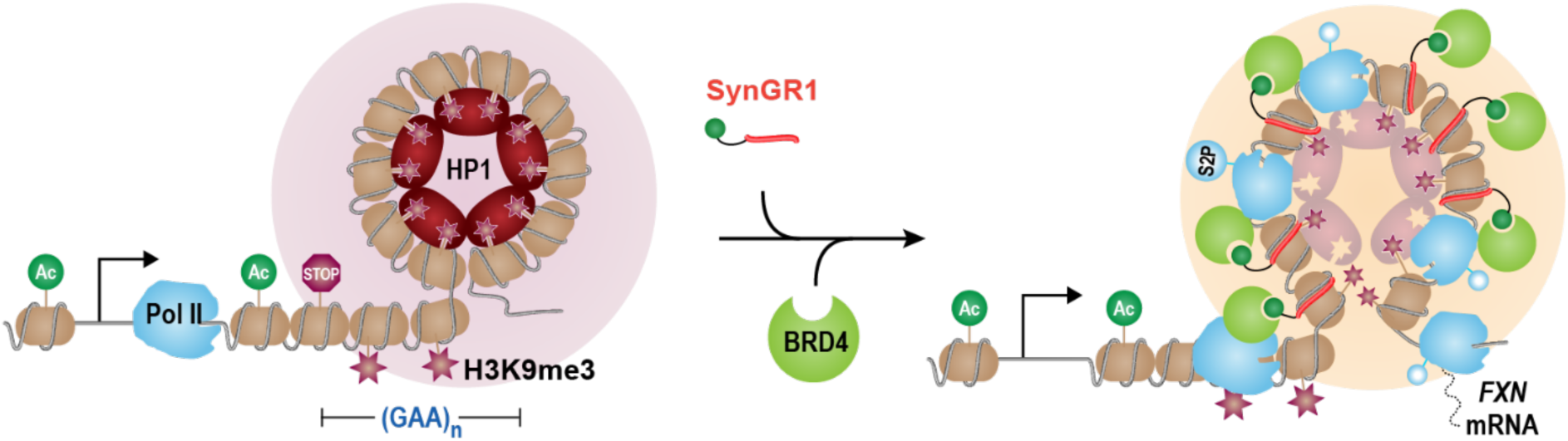
Model of transcription within the context of repressive chromatin. Repressive H3K9me3 marks (red stars) and their reader HP1 proteins (red ovals) that compact and condense chromatin (pink sphere) can function as physical and energetic roadblocks to RNA polymerase II (Pol II) dependent transcription. Acetylation of histone subunits (Ac) is a mark of active transcription. In Friedreich’s ataxia, pathological hyper-expansion of GAA trinucleotide repeats within frataxin (FXN) results in enrichment of H3K9me3 and HP1 and the consequent attenuation of transcription elongation. The precision-tailored heterobifunctional synthetic gene regulator (SynGR1) can partition into and via its DNA binding module (red bar) it can bind GAA-rich sites within this repressive chromatin. Through the chemically appended JQ1 moiety (green ball), it can bind BRD4/BET protein and leveraging the principle of “induced proximity”, SynGR1 partitions BRD4 into the HP1/H3K9me3-rich repressive chromatin without dispersal of the repressive marks or readers. This orthogonal state enables Pol II to enter and transcribe through the repressive chromatin state. The process of transcription and the generation of GAA-rich transcripts results in further deposition of repressive marks and HP1. Nevertheless, SynGR1 is able to override this dynamic and active placement of repressive marks. Beyond *FXN* and independent of SynGR1, BRD4 can access HP1α puncta in live cells and transcription within heterochromatin has been reported. BRD4/BET-enbabled expression of *FXN* provides a mechanism by which transcription could occur at other dynamically repressed genomic loci.

From the therapeutic perspective, our data indicate that “epigenetic drugs” that inhibit chromatin-modifying enzymes will perturb promoter gating more readily than the GAA-dependent gating of elongation that we observe at *FXN*. In contrast, SynGR1 is designed to remove the elongation block placed by the GAA-repeat expansions in *FXN*. The synergy gained by combining therapeutic agents that act at sequential mechanistic steps in gene transcription will expand the therapeutic index and potentially confer therapeutic benefit for FRDA patients.

## Materials and Methods

### Cell Culture and Treatment

GM15850, GM15851, GM16209, GM04079 and GM16197 cells were obtained from the Coriell Institute and cultured under recommended conditions. Cells were cultured in RPMI (GIBCO) containing 15% FBS (GIBCO) and an antibiotic/antimycotic containing 100 U/mL penicillin, 100 U/mL streptomycin, and 0.25 U/mL Amphotericin B (GIBCO). Small molecules were dissolved in DMSO and were added to fresh culture in media.

### Peptide Synthesis and Protein Production

The H3 peptide sequence ARTKQTARKSTGGKAPRKQLATKA labeled with an N-terminal 5-FAM was produced at the St. Jude Children’s Research Hospital Peptide Production Facility using a Symphony X solid phase peptide synthesizer. The lysine and serine modifications were introduced during peptide synthesis using pre-modified amino acids. Peptide purity was validated using 220 nm and 492 nm HPLC and verified using mass spectrometry.

HP1α, HP1β, and HP1γ proteins were generated by the St. Jude Children’s Research Hospital Protein Production Facility. Briefly, BL21(DE3) Rosetta *E. coli* cells were transformed with pBH4-6His-TEV-HP1α plasmid and selected using carbenicillin and kanamycin. Single colonies were grown and induced with IPTG. Pellets were lysed in lysis buffer (300 mM NaCl, 1xPBS, 10% glycerol, 7.5 mM imidazole pH 8) in the presence of PMSF, pepstatin A, aprotinin, and leupeptin. Supernatant from lysate was incubated with cobalt resin for at least 1 hour and then washed in a gravity column. Protein was eluted in 20 mM HEPES pH 7.5, 150 mM KCl, 400 mM imidazole pH 7.5. 50 μL TEV protease at 2 mg/mL was used to cleave the tag, and the protein was then dialyzed overnight in 20 mM HEPES pH 7.2, 75 mM KCl, and 1 mM DTT. The protein was then passed through a MonoQ column using standard protocols and eluted with a 200 mM-1 M KCl gradient and collected. Protein was then passed through a 0.22 μm centrifugal filter and purified further on an S75 size exclusion column. Protein was concentrated using 10 kDa spin column concentrators as needed, and concentrations were verified using UV absorbance at 280 nm (Nanodrop).

A synthetic gene for BRD4s, codon-optimized for *E. coli*, was ordered from Genscript in a pET28a vector. A hexa-histidine tag followed by a TEV protease cleavage site was placed at the N-terminus of the open reading frame. The BRD4s plasmid was transformed into BL21-RIPL cells, cultures were grown in LB medium, and expression initiated by addition of IPTG. Cells were lysed in 30 mM imidazole pH 7.8, 1 M NaCl by sonication. The clarified lysate was loaded onto a 5 mL Fast Flow Chelating Sepharose gravity column and washed with the resuspension buffer. Protein was eluted in 300 mM imidazole pH 7.8, 300 mM NaCl and subsequently diluted three-fold before loading onto a HiTrap Heparin column for the removal of bound nucleic acid. The protein was eluted with a gradient of NaCl in 20 mM HEPES pH 7.5 and concentrated to 100 μM. Protein was then dialyzed into 20 mM Tris pH 7.8, 150 mM NaCl, 5 mM DTT and flash frozen and stored at −80 °C.

### Fluorescence Polarization Binding Assays

Fluorescence polarization binding assays were performed using 20 μL FP buffer (0.02% NP-40, 150 mM KCl, 20 mM HEPES pH 7.5, 1 mM DTT) in black flat bottom plates (Corning 3821BC). HP1 proteins (20 mM HEPES pH 7.5, 200 mM KCl, 10% glycerol, 2 mM DTT) were serially diluted into the plate. 30 nM final concentration of H3 peptides were added to each well with an Echo Liquid Handler. Plates were agitated on shaker for 5 minutes and incubated at RT for 90 minutes. Fluorescence polarization of all samples was determined using fluorescence excitation at 485nm and emission at 520nm.

### Chromatin Immunoprecipitation

2.5×10^7^ cells were used for each ChIP sample. Covaris truChIP Chromatin Shearing Kits were used for fixation and nuclear isolation using the standard protocol. Immunoprecipitation was performed as follows: 35 µL/sample of Protein A/G (Pierce 88803) beads were washed 2x with and resuspended in ChIP Lysis Buffer (50 mM Tris-HCl, 10 mM EDTA, 0.5% Empigen BB, 1% SDS) and used to preclear samples for 1 hr. 8 µL of sample was added to 292µL Elution Buffer (1% SDS, 100 mM NaHCO3) and stored as input at −20°C. 800 µL of sample was added to 2 mL of IP Buffer (2 mM EDTA, 150 mM NaCL, 20 mM Tris-HCL pH 8.0, 1% Triton X-100). 6µg antibody was added to each sample, and they were rotated at 4°C overnight. Antibodies used can be found in table S1. 100 µL magnetic beads per sample were washed 2x with equal volumes of IP buffer and then added to each sample and samples incubated for 1 hour. Samples were centrifuged at 13k RPM for 1 minute and then placed on magnet. Supernatant was discarded and beads were washed with 1 mL each of Wash Buffer 1 (2 mM EDTA, 20 mM Tris-HCl pH 8.0), 0.1% SDS, 1% Triton X-100, 150 mM NaCl), Wash Buffer 2 (2mM EDTA, 20 mM Tris-HCl pH 8.0, 0.1% SDS, 1% Triton X-100, 500 mM NaCl), Wash Buffer 3 (1 mM EDTA, 10 mM Tris-HCl pH 8.0, 250 mM LiCl, 1% Deoxycholate, 1% NP-40), and 2x with 1 mL TE Buffer (10 mM Tris-HCl, 1 mM EDTA). Samples were centrifuged at 13k RPM for 30s and residual TE buffer was removed. Magnetic beads were resuspended in 300 µL Elution Buffer. 12 µL 5 M NaCl was added to both samples and inputs and placed at 65°C for 18 hours. Samples were then spun at max speed at RT and supernatant was transferred to new tubes. RNase A (Fisher EN0531) was added at 0.2 µg/µL and samples incubated at 37 °C for 1 hour. Proteinase K (Fisher 25530049) was added at 0.2 ug/µL and samples incubated at 55 °C for 1 hour. DNA was isolated using Qiagen DNA cleanup buffers PB (19066) and PE (19065). Briefly 1.2 mL buffer PB was added to each sample and samples were added to DNA columns (Fisher NC0066803) and centrifuged for 60 seconds. Columns were washed 2x with 0.75 mL buffer PE and eluted in 50 µL nuclease-free water. Samples were sequenced using Next Generation Sequencing by NovaSeq6000 by the St. Jude Children’s Research Hospital Genome Sequencing Facility.

### ChIP-seq Data Analysis

All ChIP-seq data were sequenced for this study and deposited under accession number GSE196619 in the GEO database. Collection of all sequencing reads were processed with the Trim Galore tool (available on-line at https://www.bioinformatics.babraham.ac.uk/projects/trim_galore/), removing all potential adapter sequences and quality trimming reads with cutadapt using Q20 quality score cutoff^51^. Next, reads were aligned to the human reference genome GRCh38.p12 using bwa (v0.7.17-r1198)^52^ and the output was converted to BAM format with samtools (v1.2)^53^, followed by identification of duplicated reads with bamsormadup tool from biobambam2 program (v2.0.87)^54^. Subsequently, the SPP tool (v1.11)^55^ was used to estimate fragment size with the relative strand cross-correlation analysis; and uniquely mapped reads were extracted from BAM files with samtools and extended with bedtools (v2.24.0)^56^, using the fragment size value precalculated with cross-correlation analysis. Subsequently, the MACS2 program^57^ was used to call peaks in narrow mode, with -nomodel -q 0.05 flags (high confidence peaks). In parallel, peaks were also called with more relaxed criteria, setting the -q flag to 0.5, which are here further referred to as low confidence peaks. The reproducible peaks of biological replicates were identified as those which either in both replicates had overlapping high confidence peaks or those, which in one replicate had a high confidence peak, which was supported by a low-confidence peak in the second replicate. Finally, reproducible peaks were annotated with genes if the peak overlapped the with gene promoter, defined as transcription start site (TSS) ± 2000 bp, and based on the reference annotation from Gencode^58^. For the purposes of visualization, the mapped reads’ densities were converted to BigWig format and normalized to 15 million non-duplicated mapped reads. Next, whenever applicable, the average signal between biological replicates was calculated.

Differential peak binding analysis included calculation of the fragment counts per reproducible peaks based on bedtools (v2.24.0)^56^, combined with in-house scripts, and was followed by limma-voom approach^59,60^ to differentially binding peaks. Different levels of stringency were used to classify the peaks as differentially binding, including p<0.05.

### Velocity Plots

To visualize the differences between two conditions, a new approach that utilize the vector (aka. Quiver) plots, was introduced, which for this study, covered region at coordinates chr9:69,035,000-69,039,600. These plots, which are here referred to as velocity plots, emphasize the differences of the shape and intensity of the enrichment signal between two conditions. Velocity plots are generated for a predefined genomic region of interest, and step-by-step guidelines on how the velocity plots are generated were depicted in the supplementary figure 2. The signal enrichment from two toy example conditions, both of which were pre-normalized to the same sequencing depth, and which both use the same bin size (in our case genome was separated in 50 bp long bins), are visualized on the figure 2A. The input format was bedGraph. Note that the values represented by entries of bedGraph files for each condition, which are 1-dimentional, were then converted to 2-Dimentional matrix, which dimensions depend on the number of bins that span the genomic region of interest, and the user-specified number of desired layers (in our case equal to 5 layers). Next, the algorithm identified the maximum enrichment value from both conditions, which was round up to integer. This max integer value was then divided by the number of layers, which was subsequently used to calculate m - the max enrichment value per layer. Next, for each bin, the algorithm calculated how many m values could be covered by the enrichment value from that bin, which in the final matrix were assigned the value of 1, and what was the ratio of the remaining signal to the m value (figure 2B). Finally, the values identified were ordered in ascending order and filled in the values for the layers 1-5 of the corresponding bins. On the example of 12th bin of the condition 1 (figure 2B), which enrichment value is equal to 4.3, the m value is equal to 1, so the 4.3 = 4*m + 0.3; and the 0.3/m = 0.3; therefore, the values placed in layers 1-5 were 0.3, 1, 1, 1, 1, as visualized on the figure 2B. Next, in order to compare the condition 1 to condition 2, the 2-D matrix of the condition 2 has to be subtracted from 2-D matrix of the condition 1 (figure 2C), and vice versa, to compare condition 2 to condition 1, 2-D matrix of the latter has to be subtracted from the 2-D matrix of the former. Next, the algorithm calculated the values required to plot regular quiver plot from matplotlib Python package (description available on-line at: https://matplotlib.org/stable/api/_as_gen/matplotlib.pyplot.quiver.html), that is X and Y coordinates, which correspond with the Bins and layers from the processed 2-D subtraction matrix. Moreover, the quiver plotting function requires the values of the horizontal and vertical arrow tilt to position the vector on the plot. Both were calculated by traverse through the 2-D subtraction matrix, with calculating the tilts using the weight pattern shown on figure 2D. I.e. in order to calculate the vertical arrow tilt, the value from current cell, as well as the cell corresponding to the previous layer, were taken under consideration (result values for vertical arrow tilt are shown on figure 2F); however, to calculate the horizontal arrow tilt, all the adjacent layers’ values were taken under consideration, with the weights decreasing the further the distance from the cell of origin (for the example matrix shown in figure 2C, the horizontal tilt values are shown in figure 2E). Finally, the values for the vertical arrow tilt, which range between -1 and 1, were also used to assign color from the blue-white-red color map. The resulting vector-enrichment plots for the toy example, for both the condition 1 over condition 2, and condition 2 over condition 1, are shown on the Extended Data figure 2G.

### siRNA, RNA Isolation, and qPCR

On-target plus siRNA was purchased from Dharmacon and Dharmafect I was used for siRNA transfection and knockdown using the recommended protocol. RNA was isolated using Qiagen RNeasy isolation kit (74106) and eluted in 50µL nuclease free water. RNA concentration was determined using nanodrop and diluted to 100ng/uL in nuclease free water. 100ng of RNA was used to generate cDNA using iScript reverse transcription cDNA synthesis kit (Bio-Rad 1708891). The resulting cDNA concentration was determined using nanodrop and diluted to 100ng/uL. Quantitative polymerase chain reaction was performed using SYBR Green SsoAdvanced (Bio-Rad 1725275) with 100ng cDNA per sample. qPCR was performed on ABI7900.

### RNA-seq data analysis

Raw RNA-seq reads were quality filtered and trimmed with Trim Galore tool (available on-line at https://www.bioinformatics.babraham.ac.uk/projects/trim_galore/). Reads were then aligned to the human reference genome (hg38 / GRCh38.p12) using STAR^61^. Subsequently, based on the reference annotation from Gencode (Release 31)^58^, read counts per gene were calculated using RSEM^62^, including only level 1 and 2 protein-coding genes, with at least 10 reads per condition. Next, to identify differentially expressed genes (DEGs), the remaining genes were processed using limma-voom approach^59,60^. Based on the values of log2(fold-change), p-value and false discovery rate (FDR), genes were then classified into various categories of differential expression representing different levels of stringency: |log2(FC)| > 1 and FDR < 0.05.

### Imaging

Cell culture: 18×18 #1.5 microscope coverslips (Fisher brand:12541013CA) were acid washed and stored in 95% ethanol until use. Coverslips were placed in a 6 well dish and the ethanol allowed to evaporate while sterilized by UV light and washed with PBS. Poly-D-lysine (Gibco: A38904-01) was added to the coverslips according to manufacturer’s directions before the lymphocytes—GM15851 and GM15850—were added. Lymphocytes were cultured in RPMI amended with 15% FBS and were grown for 24 hours before treatment. Cells were treated with either 1 uM SynGR1 or DMSO for an overall 0.1% DMSO. The cells were incubated for 24 hours before the RNAScope+IF procedure. See Table 1 for reagents.

**Table 1:** RNAScope+IF reagents.

| Name | Company | Reference # |
| --- | --- | --- |
| RNAScope H<sub>2</sub>O<sub>2</sub> and Protease Reagents | Advanced Cell Diagnostics | 322381 |
| RNAScope Multiplex Fluorescent Detection Reagents v2 | Advanced Cell Diagnostics | 323110 |
| RNAScope Wash Buffer Reagents | Advanced Cell Diagnostics | 310091 |
| Hs-POLR2A | Advanced Cell Diagnostics | 310451-C2 |
| Hs-FXN | Advanced Cell Diagnostics | 428961 |
| Opal 570 | Akoya Biosciences | FP1488001KT |
| Opal 620 | Akoya Biosciences | FP1495001KT |
| BRD4-alexa647 | Abcam | Ab197608 |
| HP1α -alexa488 | Abcam | Ab185018 |
| 20x SSC buffer | Invitrogen | AM9763 |
| Nuclease Free Water | Ambion | AM9932 |
| 1x PBS | Fisher BioReagents | BP24384 |

RNAScope+ Immunofluorescence: RNAScope^63^ multiplexed with immunofluorescence was performed largely by the manufacturer’s directions with one exception—the immunofluorescence and DAPI staining occurred after the RNAScope procedure. Opal dyes were diluted 1:2000. *POLR2A* was chosen as the control to ensure RNAScope was successful in all conditions imaged since *FXN* is a low expression target.

Microscope and image acquisition: Images were captured using a Zeiss 980 Airyscan 2 microscope equipped with a GaAsP-PMT detector. The Plan-Apochromat 63x/1.40 NA oil DIC M27 objective was used with a 1 AU pinhole. Channels were sequentially imaged with a bidirectional scan and 0.2 um z-step. Image dimensions and depth: 16 bit, 0.0071 um x 0.071 um x 0.200 um. Image acquisitions conditions were optimized from the WT SynGR1 treated condition. Table 2 lists microscope settings for each channel; the excitation and emission of each fluorophore was based off Zeiss’s filter assistant. Images were acquired overnight using the Zeiss multipoint scan and the software’s autofocus set to use the DAPI channel as the reference. A 5 µm window used a total of 2.5 µm above and below the midpoint of the cells in frame was used for the z-range.

**Table 2:**
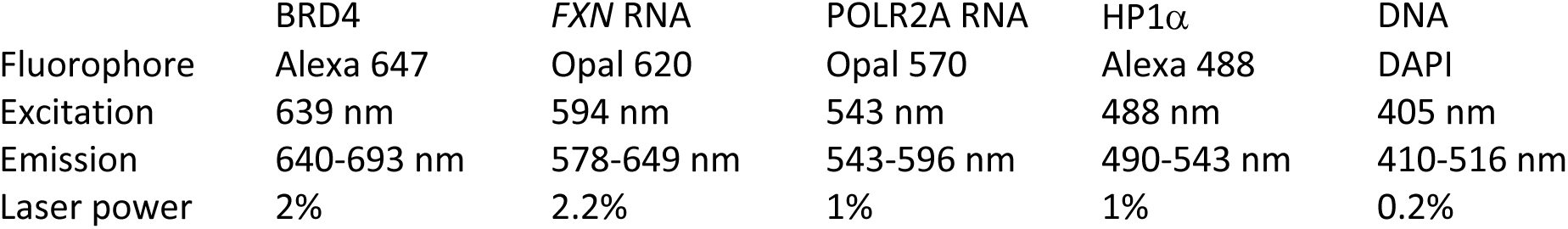
Microscope Settings for each channel.

| | BRD4 | *FXN* RNA | POLR2A RNA | HP1α | DNA |
| --- | --- | --- | --- | --- | --- |
| Fluorophore | Alexa 647 | Opal 620 | Opal 570 | Alexa 488 | DAPI |
| Excitation | 639 nm | 594 nm | 543 nm | 488 nm | 405 nm |
| Emission | 640-693 nm | 578-649 nm | 543-596 nm | 490-543 nm | 410-516 nm |
| Laser power | 2% | 2.2% | 1% | 1% | 0.2% |

### Automated 3D image analysis

Batch analysis was performed using FIJI (v1.53m) IJ1 macros. Zeiss archive files were opened using the Bioformats plugin (https://bio-formats.readthedocs.io/en/stable/users/index.html#using-bio-formats-with-imagej-and-fiji ; The Open Microscopy Environment). 3D segmentation and 3D ROI plugins are from (https://mcib3d.frama.io/3d-suite-imagej/)^64^. Images and/or cells were excluded from processing if: the image was inadvertently duplicated, a cell was incomplete, or partial image acquisition. Code will be published on Zenodo upon acceptance, doi: 10.5281/zenodo.17592796.

#### 1. Counting RNA spots in the nucleus versus in the cytoplasm

<u>MakeMasks_SLR_zeiss.ijm</u>: Images were sequentially opened in FIJI. For each cell in each image, nuclear masks were created by applying a fixed threshold to a sum intensity projection over the z-dimension on the DAPI channel. The cytoplasm was not specifically stained but was discernable via weak, non-specific background staining. Candidate masks of cells were defined by applying a fixed threshold to a sum intensity projection over all RNA-stained channels and then also over the z-dimension. Cell masks were then “exclusive-or’ed” (XOR) with the corresponding nuclear masks to create cytoplasm masks (i.e. cell with nucleus excluded) with a corresponding ROI ID. Both the nuclear and cytoplasm masks for each image were saved. The cytoplasm mask boundaries were sometimes inaccurate (e.g. when cells touched or overlapped), and a second macro allowed the user to manually edit these masks as needed (EditMasks.ijm).

<u>EditMasks.ijm:</u> Each image was opened along with its corresponding cytoplasm mask image (output of MakeMasks.ijm) and the user may manually edit the cytoplasm mask via drawing with the pencil tool in either black (to erase) or white (to add). The cytoplasm must be 8-connected around the entire nucleus, and there must be exactly one nucleus per cytoplasm. After manual editing, composite 2D ROIs were generated from the final masks, added to the 2D ROI Manager, and saved.

<u>CountSpots_zeiss.ijm:</u> Each image was opened along with its corresponding 2D composite ROI list (output of EditMasks.ijm). For each ROI in the list (corresponding to one cell), the “3D Maxima Finder” plugin was used to localize and count the spots within the nucleus and within the cytoplasm. Maxima finding was performed according to user-defined, channel-specific parameters and the same parameters were applied across all images in a set. Maxima parameters for RNA used were xy radius 8, z radius 7, and noise 10. 3D spot coordinate lists were saved as Results .csv tables.

#### 2. Measuring the concentration of protein near RNA spots versus in the nucleus

<u>SpotsConcentration1_zeiss.ijm:</u> Each image was opened, along with its corresponding 2D composite ROI list (output of EditMasks.ijm) and spots coordinate lists corresponding to a given RNA channel (output of CountSpots1.ijm). At the xyz coordinate for each spot, a 7×7-pixel 2D ROI was drawn on the protein channel of interest and the average intensity (concentration) of the protein signal was measured. Within each nucleus, the measurements for each spot were in-turn themselves averaged together and recorded as the average intensity within the spots per nucleus. On the protein channel, the z-slice corresponding to the average z-position of all the spots within a given nucleus was used to measure the average intensity within the entire nucleus. Results were saved into a .csv table.

<u>Control measures for BRD4 and HP1 α:</u> For each identified *FXN* ATS, a second 7×7 2D ROI was randomly chosen by the user to be on the same plane as the ATS, but not the same location of the ATS. BRD4 and HP1α intensity was measured at the selected location and recorded in excel.

#### 4. 3D analyses of BRD4 and HP1α

nucROI_2.ijm: An ROI file was created with the nucleus and corresponding cell ID previously generated by the editMasks code.

<u>3D_coloc_SLR6.ijm:</u> This code creates multiple 3D masks of BRD4, HP1α, and nuclear DAPI signal. Each image is opened individually and the nucROI is used to designate each nucleus. The channels are separated, and the RNA channels are removed. The BRD4 and HP1α channels undergo rolling ball background subtraction, radius 30 pixels. BRD4, HP1α, and DAPI nuclear ROIs were duplicated for further analysis. A nuclear mask of DAPI channel was created in the 3D Manager 3D Nuclei Segmentation plugin. The HP1α puncta (higher concentration regions defined by user threshold) were detected by applying a 3D unsharp mask followed by “3D Simple Segmentation”. This mask was called the punctaMask. The punctaMask was subtracted from the nuclear mask to create the nonpunctaMask. The Coloc2 plugin was used to determine the colocalization coefficients of BRD4 and HP1α in either the HP1α punctaMask or HP1α nonpunctaMask. For each analysis, Costes’s regression was used to determine the threshold value. The puncta colocalization values and the nonpuncta colocalization values were saved as a .csv file. Last, the duplicated BRD4, HP1α, DAPI, nuclear mask, punctaMask, and nonpunctaMask images were merged and saved as a .tif.

<u>3D_puncta_SLR4:</u> This code uses the .tif output from 3D_coloc_SLR4 to measure the volume and intensity of BRD4 and HP1α in the punctaMask and nonpunctaMask. Briefly, each image is opened and added to the 3D manager. For each channel, quantif was ran to determine the volume and intensity within the different masks. The data were saved as a .csv file. If no HP1α puncta were previously detected, the code creates a “noData” .csv file.

<u>Compiling FIJI Data:</u> All FIJI generated .csv sheets were compiled into manageable .xls sheets via python scripts using the St. Jude Center for Bioimage Informatics JupyterHub. A unique 5-7 digit was assigned to each ImageJ cell ID. A Master key file was generated and contained the image batch name, ImageJ cell ID, and unique ID. Compiled excel sheets were manually examined and rearranged to facilitate import into MATLAB.

### MATLAB image data quantification

All data generated by ImageJ was analyzed in MATLAB. RNA and protein analyses were based off previously established protocol^65–68^. Statistical analyses and graph generation were performed in MATLAB and Prism. The compiled excel sheets were imported into MATLAB and ran through several codes:

<u>Protein_cell:</u> The BRD4 and HP1α protein intensity signal was normalized by subtracting the background of each image. The output was used to normalize BRD4 and HP1α intensity at the *FXN* active transcription site(s) (ATS).

<u>Process_RNA</u>: The datastat function was used to determine the median cytoplasmic RNA intensity. Candidate RNA spots were removed if the spot was outside +/- the standard deviation from the median. All RNAs were normalized to the intensity of the filtered mRNA median. The normalized intensity values were rounded to the nearest integer.

To determine the active transcription sites, nuclear RNAs that had intensity values greater than the median value of a single mRNA + 1 std (∼ 2 RNAs when adjusted for RNA count) were placed into a candidate list to be tested. All nuclear RNAs that were below this threshold were considered mRNAs and added to the mRNA list. All RNAs that rounded to zero were removed.

Fold: After background subtraction that occurred in the protein_cell code, BRD4 and HP1α intensity at the ATS were divided by the average intensity of each protein in the cell to calculate the fold change of each protein at the ATS.

### Protein Labeling

Protein was dialyzed into 10 mM HEPES pH 7.5, 150 mM NaCl, 5 mM DTT. HP1α of HP1γ was labeled with Oregon Green (Fisher O6147) and BRD4s was labeled with Rhodamine Red (Thermofisher R6160). Briefly the dye was dissolved at 100 mM in DMSO and added to protein in a 20:1 dye-to-protein molar ratio and incubated on a shaking platform for 5 hours. The reaction was quenched with 2 M excess Tris. Protein was then dialyzed into 20 mM Tris pH 7.8, 150 mM NaCl, and 5 mM DTT.

### Polyamide Synthesis

Polyamide (PA1) was synthesized using solid phase peptide synthesis^51,52^ and then conjugated to activated JF646-NHS ester by using DIPEA and DMF. The reaction was carried out at room temperature for 6h. After completion of the reaction, diluted in 15% of acetonitrile in H2O and injected in Prep-HPLC to purify the compound. The pure fractions were collected and lyophilized to obtain pure compound.

### Phase Separation Assays

DNA oligos were purchased from IDTDNA, resuspended in 50 mM NaCl and annealed at 95 °C for 5 minutes and ramped to 4 °C at 0.1 degree/s. DNA was re-annealed before each experiment and incubated with polyamide for at least 3 h at 4 °C. HP1α, HP1γ and BRD4s were incubated with DNA/polyamide in dialysis buffer for 1 hour before imaging on a Nikon C2. Time course experiments were performed by pre-forming HP1α condensates in PCR tubes for 1hr, transferring the solution to a coverslip affixed to a perforated 35 mm glass plate, and then adding BRD4s at 50 nM. The samples were imaged on the Marianas 2 and images were taken every 5 seconds. Super-resolution microscopy was performed on a Zeiss LSM980 Airyscan microscope on a solution in the 2-phase regime equilibrated for 90 minutes.

### Quantitative Analysis of Synergistic Interactions

Drug combination studies were evaluated using Bliss independence^69^. Briefly, all unique pairwise combinations of RGFP109 (DMSO, 6.25, 12.5, 25, 50) and SynGR1 (DMSO, 0.125, 0.25, 0.5, and 1 uM) were evaluated in GM15850 cells using qPCR after 24 hours of drug treatment. Gene expression values from three independent biological replicate experiments were averaged and then normalized to the range 0 (the expression induced by DMSO) to 1 (the expression induced by the combination at the highest concentration of each drug tested) to generate the observed Bliss surface. The expected Bliss surface was calculated from the behavior of each drug as single agent, and then subtracted from the observed Bliss surface to yield the differential Bliss surface. For the differential Bliss surface, values >0 indicate synergy, values = 0 are additive, and values < 0 are antagonistic.

**601 DNA FWD** CTGGAGAATCCCGGTCTGCAGGCCGCTCAATTGGTCGTAGACAGCTCTAGCACCGCTTAAACGCACGTACGCGCTGTCCCCCGCGTTTTAACCGCCAAGGGGATTACTCCCTAGTCTCCAGGCACGTGTCAGATATATACATCCTGT

**601 DNA REV** ACAGGATGTATATATCTGACACGTGCCTGGAGACTAGGGAGTAATCCCCTTGGCGGTTAAAACGCGGGGGACAGCGCGTACGTGCGTTTAAGCGGTGCTAGAGCTGTCTACGACCAATTGAGCGGCCTGCAGACCGGGATTCTCCAG

**GAA DNA FWD** GAAGAAGAAGAAGAAGAAGAAGAAGAAGAAGAAGAAGAAGAAGAAGAAGAAGAAGAAGAAGAAGAAGAAGAAGAAGAAGAAGAAGAAGAAGAAGAAGAAGAAGAAGAAGAAGAAGAAGAAGAAGAAGAAGAAGAAGAAGAAGAAGAA

**GAA DNA REV** TTCTTCTTCTTCTTCTTCTTCTTCTTCTTCTTCTTCTTCTTCTTCTTCTTCTTCTTCTTCTTCTTCTTCTTCTTCTTCTTCTTCTTCTTCTTCTTCTTCTTCTTCTTCTTCTTCTTCTTCTTCTTCTTCTTCTTCTTCTTCTTCTTC

**Table S1.**
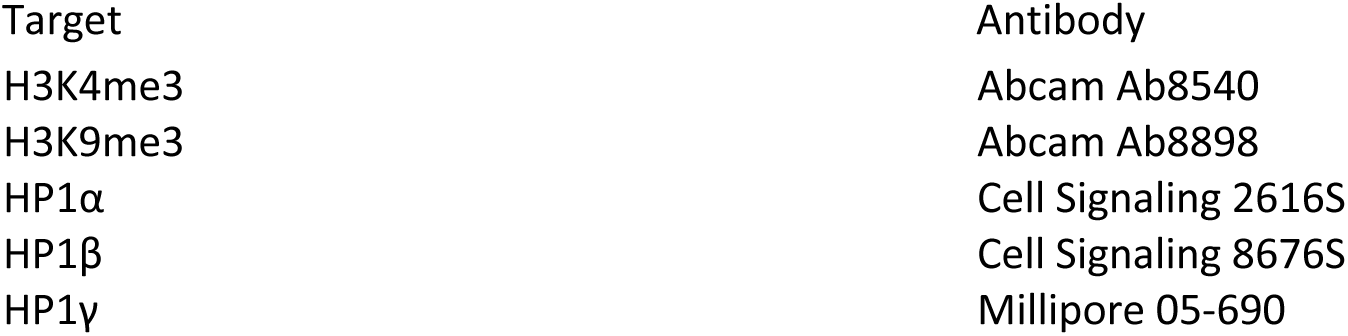

**Table S2.**
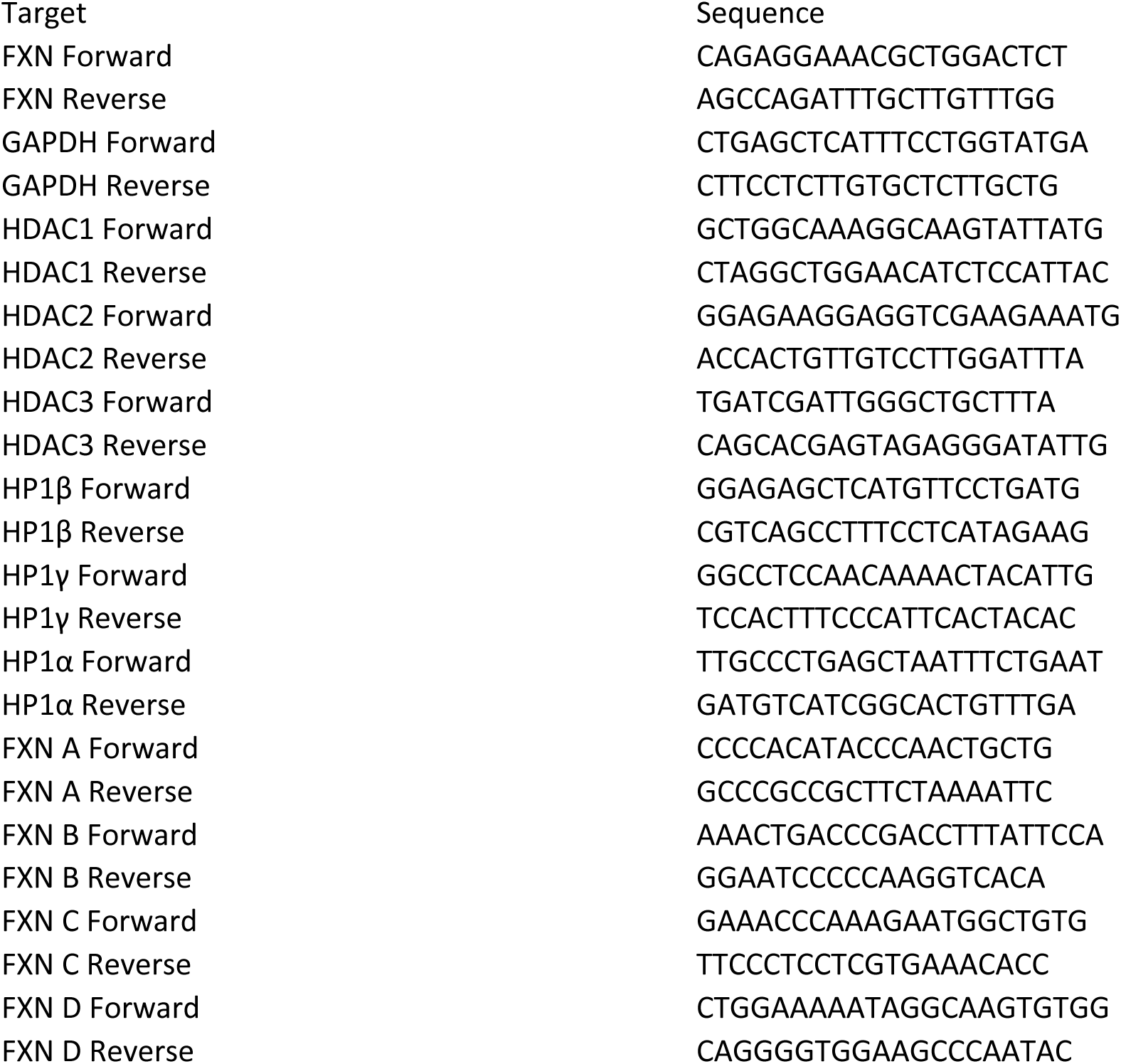

### Compound Screening and global siRNA knockdown

GM04078 fibroblasts were grown in Minimum Essential Medium with Earle’s Salts (Gibco 41200-038) supplemented with 15% FBS (Biowest S1620, Hyclone SH30396.03), Non-Essential Amino Acids (GenClone 25-536), Penicillin/Streptomycin (Gibco 151140-122), and GlutaMAX (Gibco 35050061) at 37° C and 5% CO_2_. Cell growth and density were monitored by phase microscopy and Trypan Blue exclusion (Gibco 15250061) using a Countess 3 (Thermo Scientific A49862).

Compound Screening:

Compounds were first tested at a final concentration of 10 µM from 10 mM stocks in 100% DMSO. 25 nl of each tested compound and an additional 25 nl of either DMSO or SynGR1 were added to each well of a 384-well CulturPlate (Revity 6007688) using a Beckman Coulter Echo 655 acoustic liquid handler. 2000 GM04078 cells in 25 µl of media were then plated into each well with a final DMSO concentration of 0.2%. Hit compounds were tested in a dose-response format in 1:3-fold dilutions curves from a top screening concentration of 10 µM. 30 nl of each tested compound and an additional 30 nl of either DMSO or SynGR1 were added to each well of a 384-well CulturPlate (Revity 6007688) using a Beckman Coulter Echo 655 acoustic liquid handler. 2000 GM04078 cells in 30 µl of media were then plated into each well using a Multidrop Combi (Thermo Scientific 5840300) with a final DMSO concentration of 0.2%.

Cells were grown for 24 hours and mRNA extracted using 384-well TurboCapture mRNA plates (Qiagen 72271) per the manufacturer’s protocol. To detect the relative levels of *FXN* and *GAPDH* in each sample 4 µl of each eluted mRNA was combined with 6 µl of Luna Universal Probe One-Step RT-qPCR mix (New England Biolabs E3006E) containing the primers and probes below.

Each sample was reverse transcribed (10 minutes at 55° C and 1 minute at 95° C) and amplified (40 cycles of 5 seconds at 95° C and 40 seconds at 60° C) using a QuantStudio 7 Pro (Thermo Scientific). All curves were analyzed using Design & Analysis (Thermo Scientific Version 2.7.0) and the relative amount of FXN to GAPDH in each sample determined using 2^-ΔΔCt^. For dose-response calculation all Design & Analysis data points were imported and analyzed for 2^-ΔΔCt^ using GeneData Screener Analyzer with 1 µM SynGR1 alone as a “Neutral Control” with 0% activity.

FXN qPCR Primers and Probes:

FXN_Forward: GAGGAAACGCTGGACTCTTTAG FXN_Reverse: CACTCCCAAAGGAGACATCATAG

FXN-Probe: /56-FAM/ACCTTGCAG/ZEN/ACAAGCCATACACGT/3IABkFQ/ GAPDH_Forward: GGTGTGAACCATGAGAAGTATGA

GAPDH_Reverse: GAGTCCTTCCACGATACCAAAG

GAPDH_Probe: /5SUN/AGATCATCA/ZEN/GCAATGCCTCCTGCA/3IABkFQ/

siRNA Screening:

siRNA pools (Horizon OnTarget Plus) to each gene of interest were tested at a final concentration of 22.5 nM from 5 µM stocks in RNA resuspension buffer (Horizon B-002000-UB-100). 112.5 nl of a tested siRNA was added to one well of a 384-well CulturPlate (Revity 6007688) using a Beckman Coulter Echo 655 acoustic liquid handler. 10 µl of Opti-Mem (Thermo Scientific 51985034) containing 1% Dharmafect 1 (Horizon Discovery T-2006-01) was added to each well for 20 minutes using a Multidrop Combi (Thermo Scientific 5840300). 15 µl of complete media containing 2000 GM04078 cells was then added to each well and incubated overnight. The following day an additional 25 µl of complete media was added to each well and the plates were incubated for an additional 48 hours. A final addition of 10 µl of complete media containing 6 µM SynGR1 to a 1 µM final concentration or DMSO was added to each well for 24 hours. mRNA extraction and qPCR were then performed as above.

### Hierarchical network model

The activity based hierarchical network graph was visualized in Cytoscape (v.3.4.0) using the yFiles circular layout. Node-edge relationships were defined by hierarchically decomposing the reported targets of the epigenetic modulator using annotations from the IUPHAR/BPS Guide to Pharmacology^70^ or manual annotation when required. Compounds with multiple targets are represented by distinct leaf nodes.

Chemical space analysis was conducted using KNIME (v.5.3.3) and the RDKit^71^ radius 3 morgan fingerprint set to 1024 bits and the following t-SNE^72^ parameters: dimensions = 2, interatations = 5000, theta = 0.2, perplexity = 30.0, and threads= 20.

## Extended Data figures

**Extended Data Figure 1.**
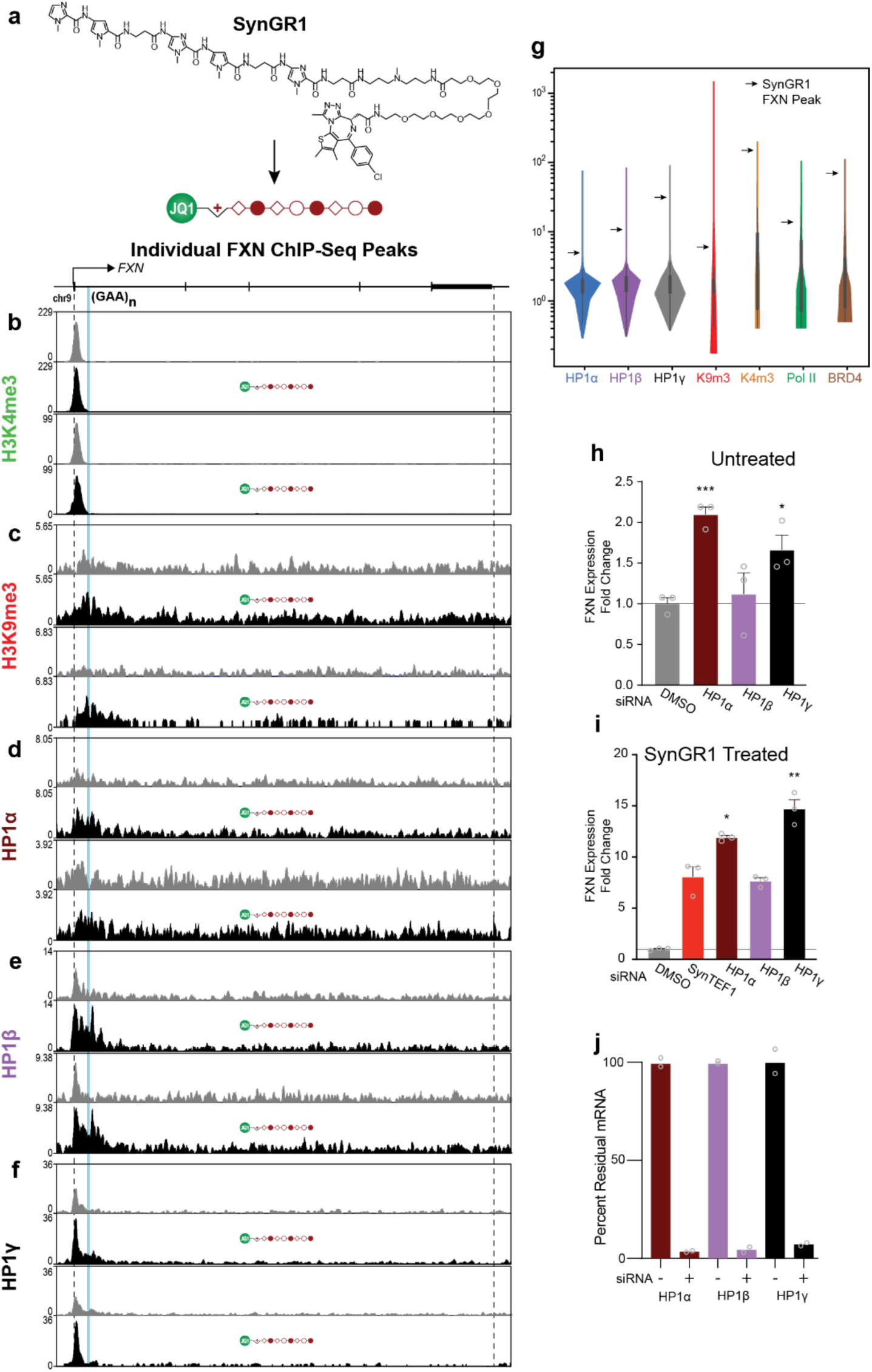
Epigenetic silencing caused by (GAA)n repeat expansion at FXN remains after SynGR1 mediated transcriptional activation. **(A)** Chemical structure of SynGR1/SynGR1. **(B-E)**, Individual ChIP-seq tracks for H3K4me3 **(B)**, H3K9me3 **(C)**, HP1α **(D)**, HP1β **(E)**, and HP1γ **(F)** treated with DMSO or SynGR1. **(G)**, Violin plot of vehicle treated ChIP-seq peaks. Arrows show FXN peaks after SynGR1 treatment. **(H)**, FXN expression for HP1 proteins after siRNA treatment in GM15850 FRDA lymphoblasts. Expression normalized to DMSO. Error bars are SEM, p-values measured by unpaired t-test compared to DMSO. HP1α p<0.001, HP1γ p<0.05 **(I)**, FXN expression after siRNA knockdown and SynGR1 treatment normalized to DMSO. Error bars are SEM, p-values measured by unpaired t-test compared to SynGR1. HP1α p<0.05, HP1γ p<0.01. **(J)**, Percent remaining mRNA after siRNA treatment compared to untreated cells.

**Extended Data Figure 2.**
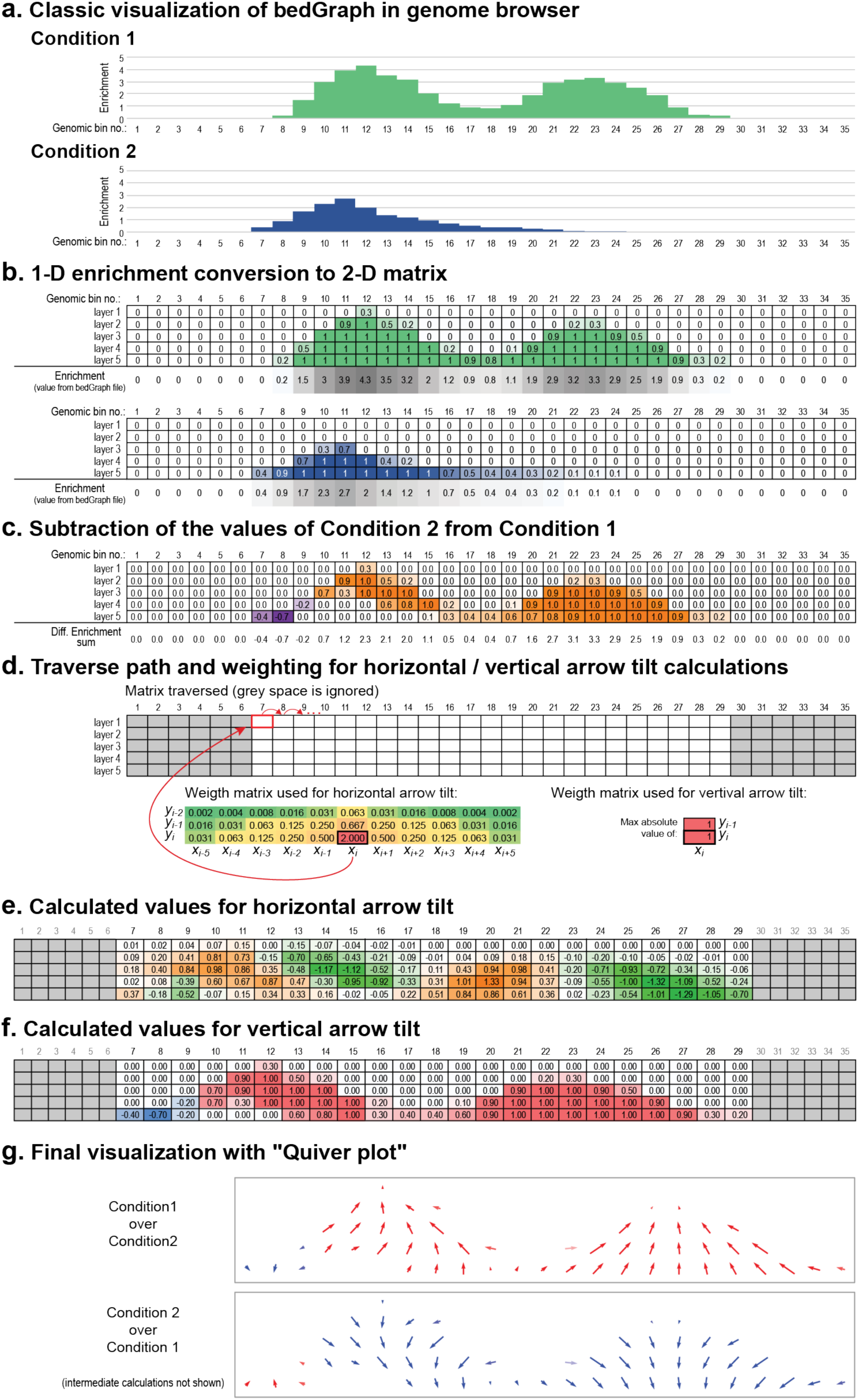
Velocity plots are a method to visualize the change in ChIP-seq peak distribution. **(A)**, Signal enrichment from two toy example conditions, both of which were pre-normalized to the same sequencing depth, and which both use the same bin size (in our case genome was separated in 50 bp long bins), are visualized. **(B)**, Layering of the max integer peak value. **(C)**, Graphical representation of the 2D matrix difference. Positive values are in orange and negative values in purple. **(D)**, Horizontal and vertical weight matrix to calculate arrow tilt. e-f, Horizontal **(E)** and vertical **(F)** weight matrix calculations for example peaks. **(G)**, Veloctiy plots generated comparing test condition 1 to 2 or condition 2 to 1.

**Extended Data Figure 3.**
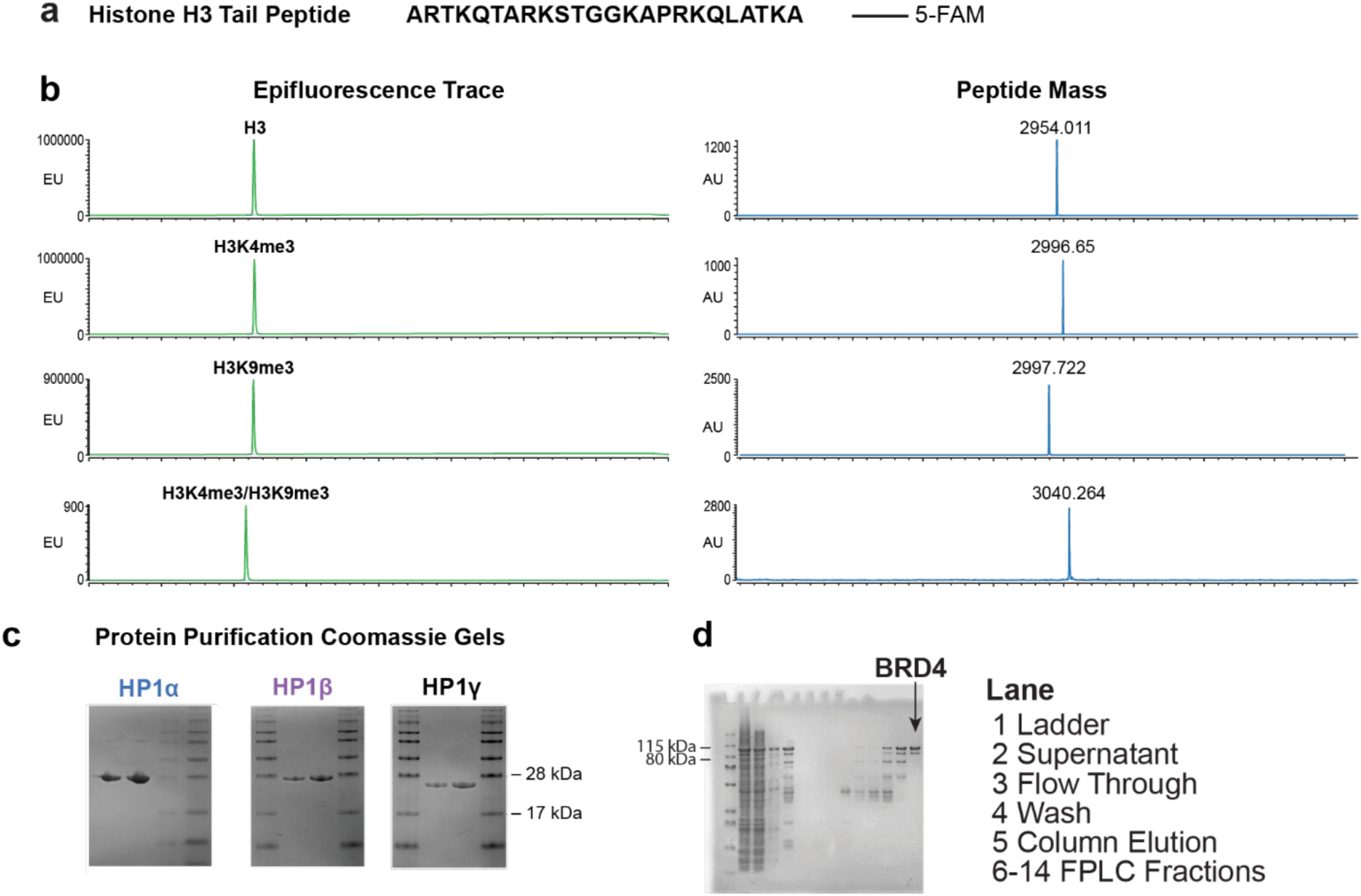
Labeled H3 modified peptide and protein generation. (A), Figure showing sequence of H3 tail peptide created with C-terminal 5-FAM. **(B),** HPLC epifluorescence traces and MALDI-TOF spectra of modified H3 tail peptides after purification showing purity of histone tail peptide. **(C-D),** Coomassie gels of purified recombinant **(C)**, HP1α, HP1β, and HP1γ. **(D),** BRD4

**Extended Data Figure 4.**
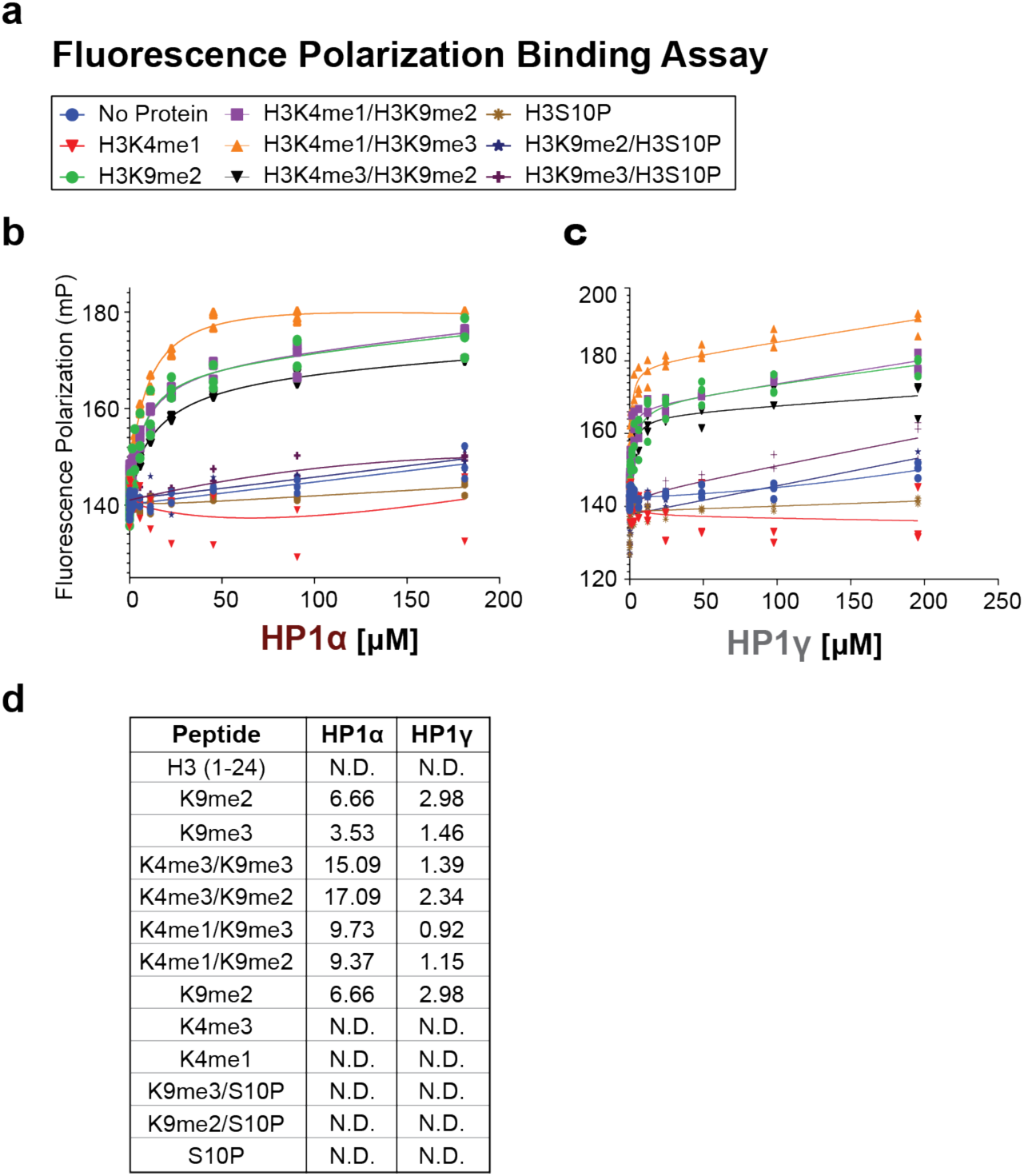
HP1 binding of modified H3 tail peptides by fluorescence polarization. (A), List of all the peptides used for Fluorescence polarization (FP) binding assay. FP binding assay of HP1α **(B)**, and HP1γ **(C)**, to all listed peptides. **(D)**, KD values of specified HP1 protein binding to modified H3 tail peptides.

**Extended Data Figure 5.**
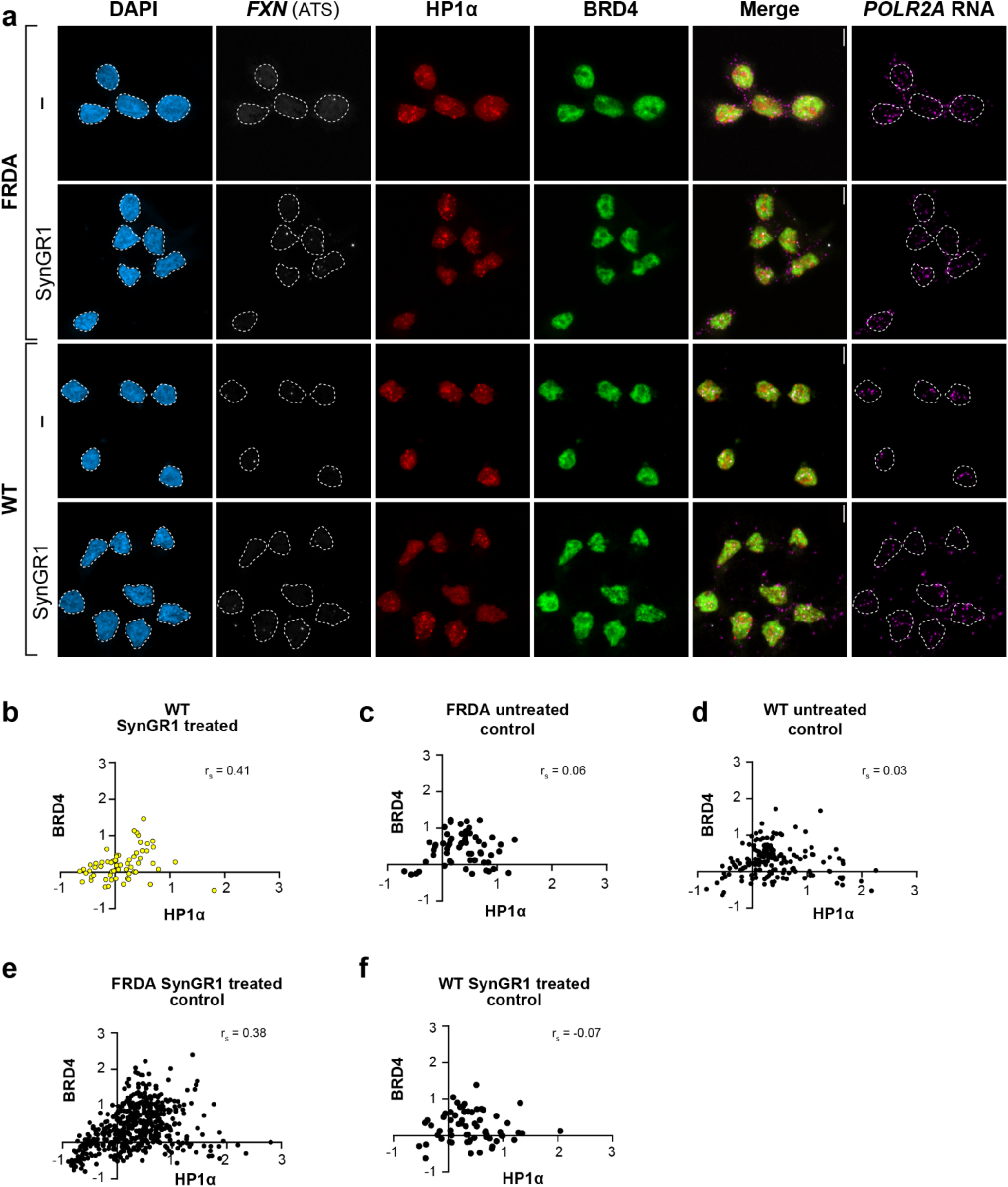
SynGR1 treatment in FRDA and WT lymphocyte cells. **(A)** Larger view of cells shown in Figure 2a. z- maximum projection shown; white dashed line--nucleus. Scale, 5 μm. **(B-F),** BRD4 and HP1α enrichment at the **(B)** FXN ATS and **(C-F)** control ROI’s of the same size as an FXN ATS for all experimental conditions. rs, spearman’s correlation.

**Extended Figure 6.**
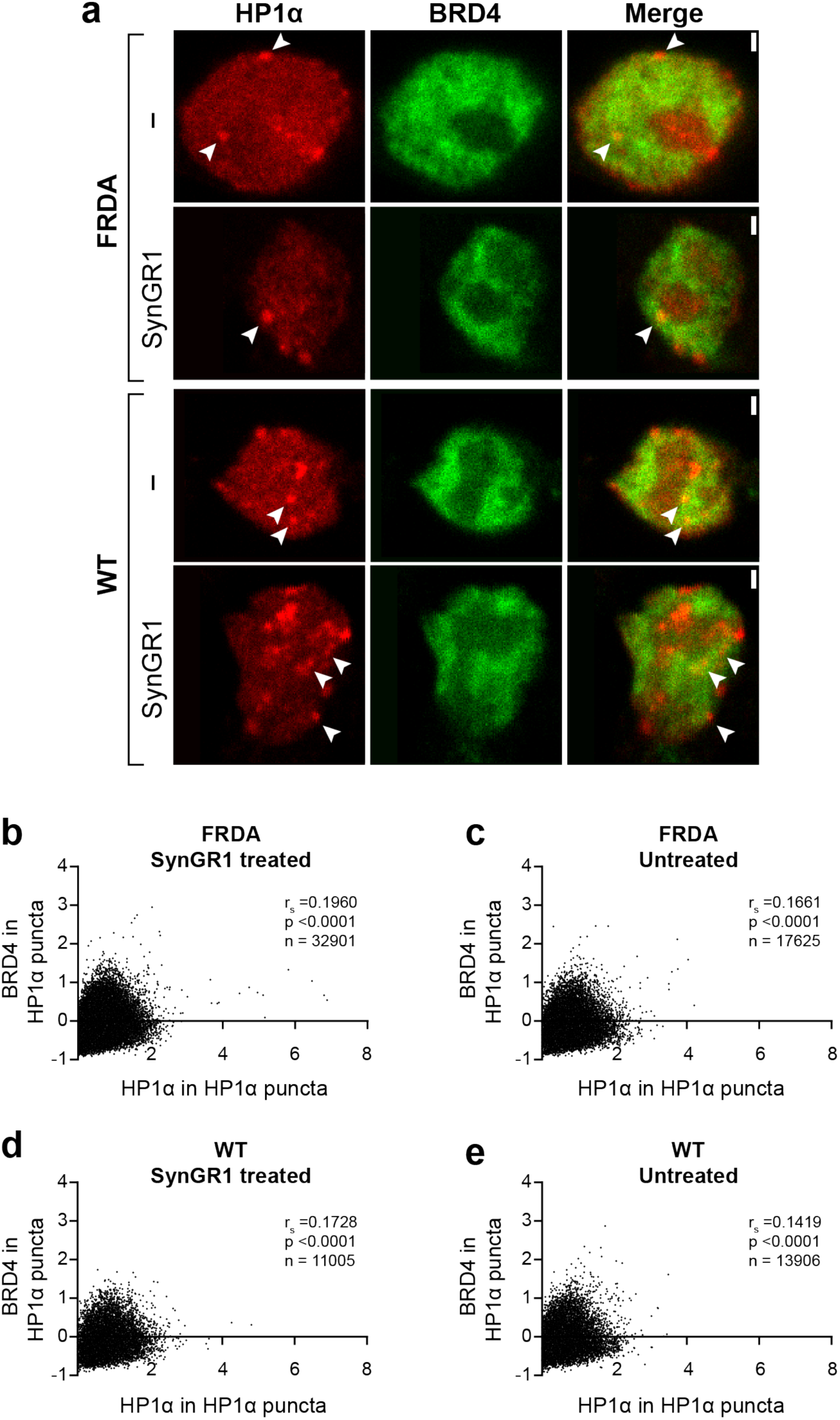
BRD4 and HP1a co-occurrence in cells. **(A)** Single z-plane images of BRD and HP1α in treated and untreated FRDA and WT cells. Arrow, regions of BRD4 and HP1α co-occurrence. **(B-E)** Normalized BRD4 fold change in normalized HP1α puncta. Briefly, a 3D nuclear mask was made for each image. For each nucleus, 3D HP1α puncta were detected. The HP1α puncta were removed from the 3D masks to get the nuclear, non-HP1α puncta region of the nucleus. BRD4 and HP1α intensity and volume was measured in the HP1α puncta and nuclear non-HP1α puncta region. For the HP1α and nuclear non-HP1α puncta region, HP1α and BRD4 concentration was calculated by dividing the intensity by the volume. To normalize the concentration in the HP1α puncta, the concentration of either BRD4 or HP1α was divided by the nuclear non-HP1α puncta concentration for each protein. The concentration was then rescaled so that depletion would be negative and enrichment positive.

**Extended Data Figure 7.**
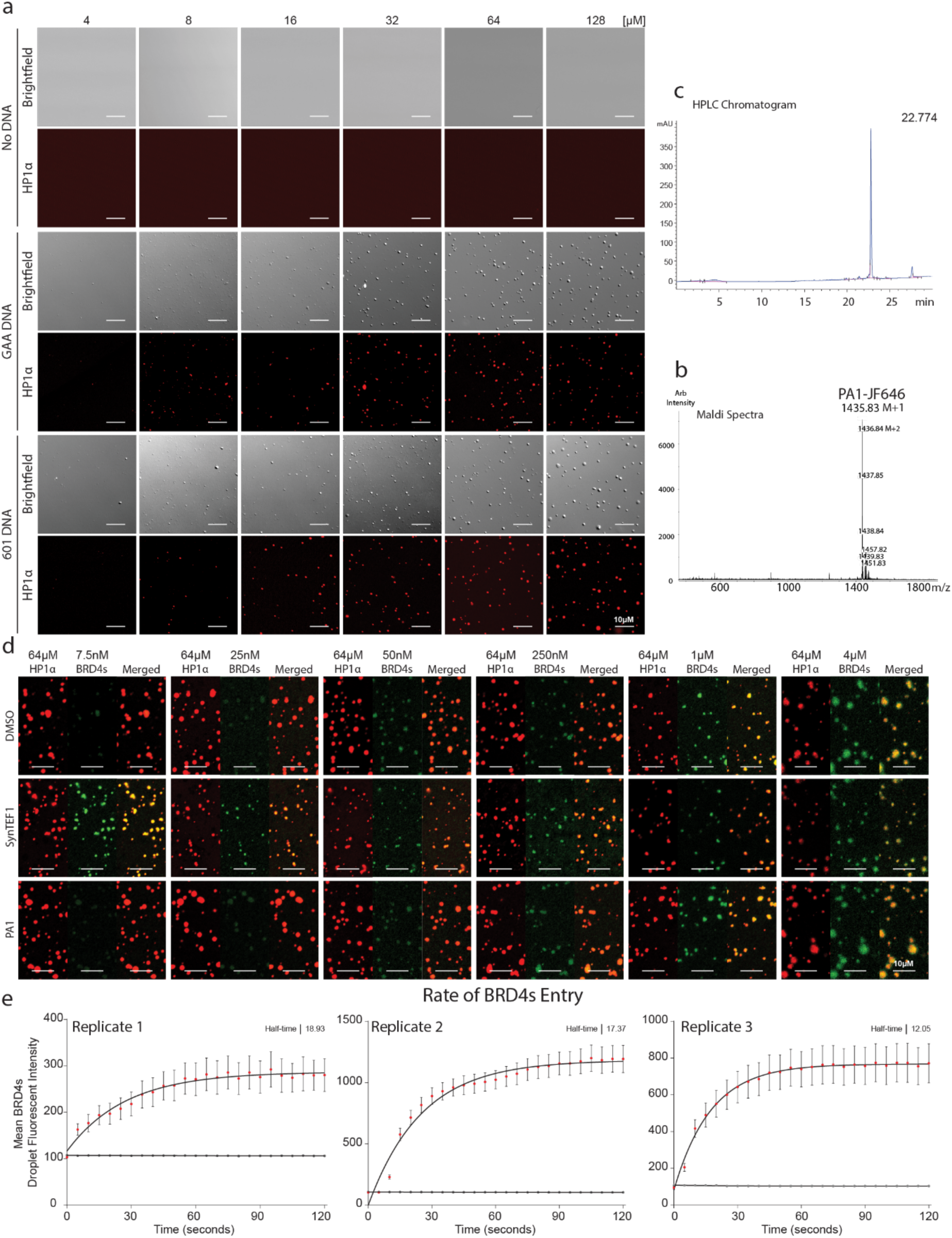
BRD4 is recruited to HP1α phase separated condensates by SynGR1. a, Titration of HP1α without DNA, with 601 DNA, or with GAA DNA showing increased phase separation with increasing protein concentration. b, MALDI TOF spectra of PA1-JF646 showing purity. c, HPLC Chromatogram of PA1-JF646 after purification. d, Titration of BRD4s with 64μM HP1α either in the presence of DMSO vehicle, SynGR1, or PA1. e, Individual replicates of timed entry of BRD4s into HP1α preformed condensates. SEM between droplets in each replicate is shown. Half-time calculated by one-phase association.

**Extended Data Figure 8.**
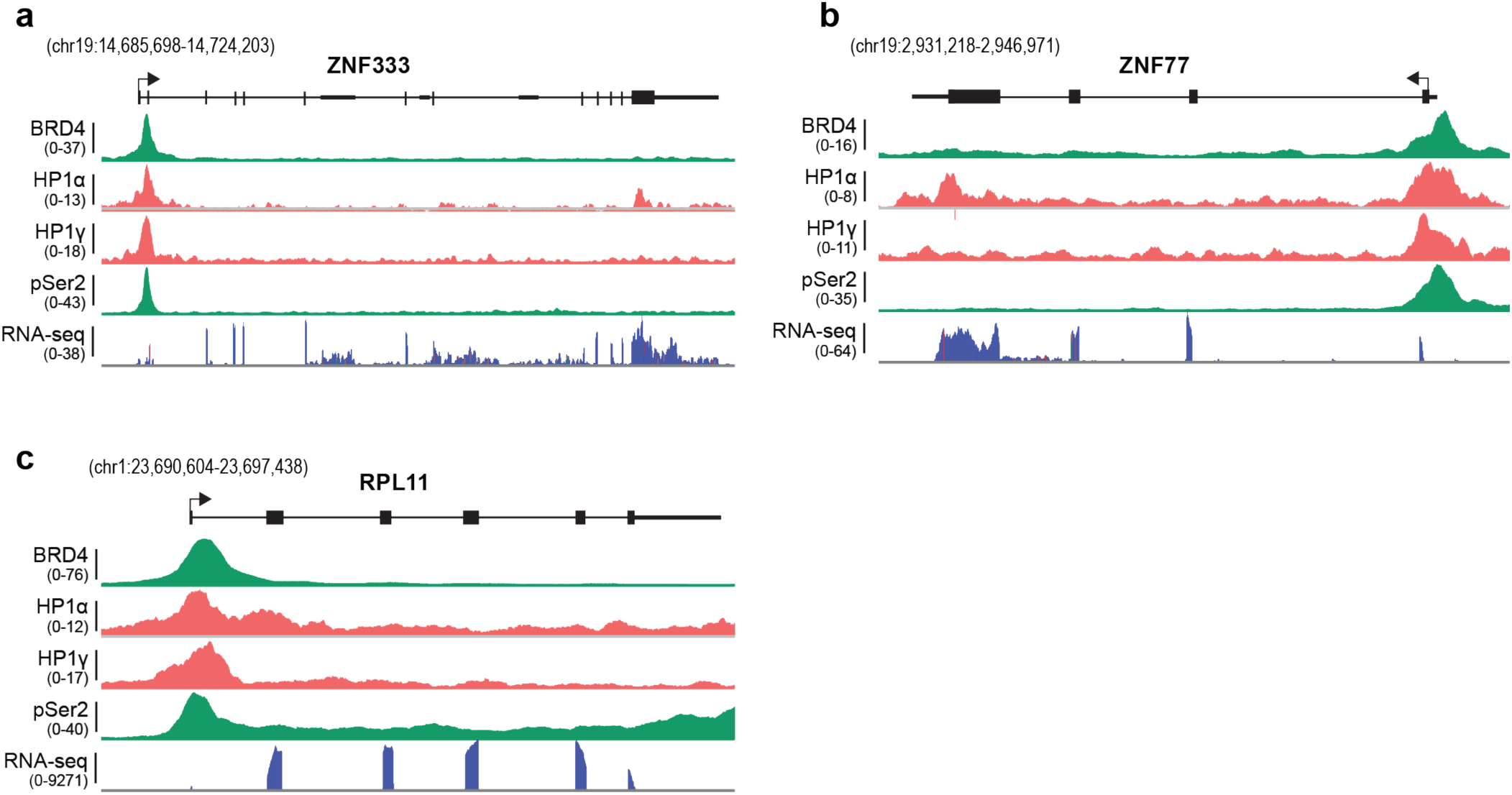
BRD4 and HP1 isoforms co-occupy active gene loci. (a–c), Genome browser views of three representative loci showing ChIP-seq profile for BRD4, HP1α, HP1γ, pSer2, together with RNA-seq. At each locus, BRD4 and HP1α/γ co-localize with Pol II over transcriptionally active regions, coinciding with RNA output. **(a)** ZNF333 gene **(b)** ZNF77 gene **(C)** RPL11 gene.

**Extended figure 9:**
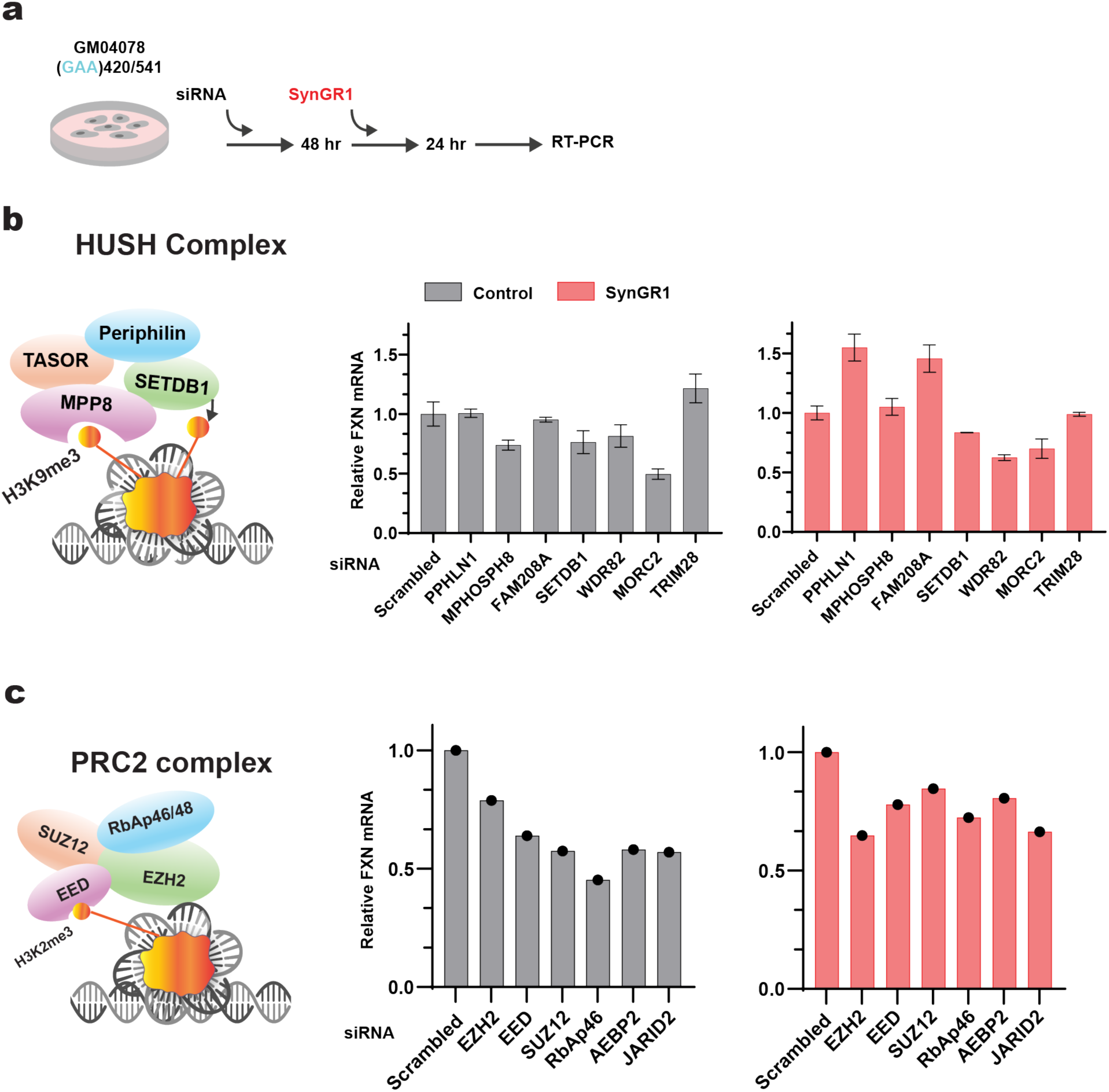
Role of PRC2 AND HUSH complexes on *FXN* gene transcription revealed by siRNA knockdown. **(A)** Experimental design showing siRNA knockdown, SynGR1 treatment, and downstream analysis. **(B-C)** SynGR1 mediated *FXN* expression after HUSH complex **(B)** or Polycomb Repressor (PRC2) complex, **(C)** knockdown. Left, composition of the respective complexes; right, *FXN* expression after knockdown of individual complex members. Gray, untreated control; Red, SynGR1 treated.

## Acknowledgements

We thank Geoff Neale and Scott Olsen of the Hartwell Center Sequencing Facility, Youming Shao and Richard Heath of the Protein Production Facility; Tomi Mori, Samira Deshpande, and Yue Pan of Biostatistics; Aaron Pitre of CTIC-LM; and Gang Wu for advice, materials, and services that enabled this study. HP1 plasmids and purification protocols were generously provided by Geeta Narlikar and Emily Wong. We thank Caitlin Deane, Ines Chen, Sweta Kumari, and members of the Ansari lab for invaluable discussions and edits, and Laura Vanderploeg for the artwork. This research included experiments conducted by the Cell and Tissue Imaging Center - Light Microscopy (CTIC-LM) and the Cytogenetic Shared Resource which is supported in part by ALSAC and the National Cancer Institute grant P30 CA021765.

## Author contributions

Author contributions: CJB, SRT, MK, WR, MC, JL, WL, TC, TM, and AZA designed research; CJB, SRT, MK, WR, MC, JB, JK, WL, JL, AM, AD, SR, MV, SK performed research; AM contributed new reagents; BY, TN, PR provided key reagents; CJB, SRT, MK, WR, MC, JB, JL, AD, SR, JO, SJK, AT, BH, KK, BX, TC, TM and AZA analyzed the data; and CJB, SRT with AZA wrote the paper.

## Materials & Correspondence

Please contact Aseem Ansari for material requests.

